# ERLIN1/2 scaffolds bridge TMUB1 and RNF170 and restrict cholesterol esterification to regulate the secretory pathway

**DOI:** 10.1101/2024.01.17.576039

**Authors:** Matteo Veronese, Sebastian Kallabis, Alexander Tobias Kaczmarek, Anushka Das, Lennart Robers, Simon Schumacher, Alessia Lofrano, Susanne Brodesser, Stefan Müller, Kay Hofmann, Marcus Krüger, Elena I. Rugarli

## Abstract

Complexes of ERLIN1 and ERLIN2 form large ring-like cup-shaped structures on the endoplasmic reticulum (ER) membrane and serve as platforms to bind cholesterol and E3-ubiquitin ligases, potentially defining functional nanodomains. Here, we show that ERLIN scaffolds mediate the interaction between the full-length isoform of TMUB1 and RNF170. We identify a luminal N-terminal conserved region in TMUB1 and RNF170 required for this interaction. Three-dimensional modelling shows that this conserved motif binds the SPFH domain of two adjacent ERLIN subunits at different interfaces. Protein variants that preclude these interactions have been previously linked to hereditary spastic paraplegia (HSP). By using omics approaches in combination with phenotypic characterisation of cells lacking both ERLINs, we demonstrate a role for ERLIN scaffolds in maintaining cholesterol levels in the ER by favouring transport to the Golgi over esterification, thereby regulating Golgi morphology and the secretory pathway.

## Introduction

The endoplasmic reticulum (ER) is the largest membrane-bound intracellular organelle and is involved in essential functions, including protein and lipid biosynthesis and calcium homeostasis. Similar to other cellular membranes, the ER lipid bilayer is organised in protein and lipid nanodomains, which define its major regions (nuclear envelope, cisternae, and peripheral tubules), compartmentalize specific processes, and mediate dynamic interactions with other cellular organelles at contact sites (English and Voeltz, 2013). E3-ubiquitin ligases embedded in the ER membrane regulate the turnover of ER-resident proteins and remove misfolded secretory cargos during biogenesis in a process called ER-associated degradation (ERAD), thus preventing proteotoxicity (Christianson and Carvalho, 2022). How these regulatory enzymes are compartmentalised in the ER membrane is poorly understood.

The homologous ERLIN1 and ERLIN2 are ER-localised members of the evolutionary conserved SPFH (stomatin/prohibitin/flotillin/HflKC) family of proteins, characterized by a conserved module of ∼180-200 amino acids, which functions to scaffold lipids and proteins (Browman et al., 2007). Topological studies of ERLIN2 have shown that it is a type-II ER membrane protein, containing a four amino acid long cytosolic tail, followed by a transmembrane domain, the SPFH domain and a coiled-coil domain (Browman et al., 2007). Like other members of the SPFH family, ERLIN1 and ERLIN2 assemble in a large ring-shaped hetero-oligomeric complex, likely formed by 24 subunits (Qiao et al., 2022; Yokoyama and Matsui, 2023). ERLIN1 and ERLIN2 have been first isolated as components of cholesterol-rich domains of the ER (Browman et al., 2006). Subsequent studies have established that ERLINs can directly bind cholesterol (Huber et al., 2013; Hulce et al., 2013). Moreover, acute downregulation of ERLIN1 and ERLIN2 leads to the canonical activation of sterol regulatory element binding proteins (SREBPs) and their target genes, leading to accumulation of lipid droplets (LDs) (Huber et al., 2013). As underlying mechanism, a role of ERLINs to bind and stabilize INSIG1 has been proposed (Huber et al., 2013).

Several evidence link ERLIN1 and ERLIN2 to ERAD, via binding and regulating membrane-embedded E3 ubiquitin ligases. ERLINs have been found in association with gp78 (AMFR) (Jo et al., 2011b), RNF170 (Lu et al., 2011; Pearce et al., 2007), RNF185 (Fenech et al., 2020; van de Weijer et al., 2020), and RNF5 (Fenech et al., 2020), although the exact organization of these complexes has not been established yet. In concert with RNF170, ERLIN2 is involved in the ERAD of the inositol 1,4,5 triphosphate (IP_3_) receptors and other model ERAD substrates. ERLIN2 associate with the IP_3_ receptor shortly after their activation and recruits the RING-domain protein RNF170 to ubiquitinate the IP_3_ receptors (Lu et al., 2011; Pearce et al., 2007). ERLIN2 and gp78 were implicated in the sterol-accelerated degradation of 3-hydroxy-3-methylglutaryl-coenzyme reductase (HMGR), a rate-limiting enzyme for cholesterol synthesis (Jo et al., 2011a). Moreover, a role of ERLIN2 in regulating the ubiquitination of the WNT secretory factor EVI was recently determined (Wolf et al., 2021). Finally, ERLINs compartmentalize the intramembrane protease RHBDL4 to control proteolysis of aggregation-prone luminal ERAD substrates (Bock et al., 2022). Altogether, these data have fostered the hypothesis that ERLINs define ER cholesterol-enriched nanodomains involved in different ERAD and proteolytic pathways.

Elucidating how ERLIN complexes interact with different E3-ubiquitin ligases and which cellular processes are controlled by the ERLIN complexes is crucial, since mutations in *ERLIN1* or *ERLIN2* gene have been associated with motoneuronal diseases, such as hereditary spastic paraplegia (HSP), primary lateral sclerosis, and amyotrophic lateral sclerosis (ALS) (Al-Saif et al., 2012; Alazami et al., 2011; Kume et al., 2021; Park et al., 2020; Qiao et al., 2022; Rydning et al., 2018; Srivastava et al., 2020; Tunca et al., 2018; Wakil et al., 2013; Yildirim et al., 2011).

Here, we show that complexes of ERLIN1 and ERLIN2 prevent cholesterol esterification, thereby promoting cholesterol trafficking from the ER to the Golgi and regulating the secretory pathway. In addition, we demonstrate that ERLIN scaffolds mediate the interaction between RNF170 and a long isoform of TMUB1 (TMUB1-L). TMUB1-L and RNF170 bind ERLIN monomers via a conserved domain, and the interaction interface in ERLIN1 and ERLIN2 is targeted by pathogenic mutations in HSP. Our data define the ERLIN complex as a crucial scaffold that couples cholesterol homeostasis to regulatory ERAD pathways controlled by RNF170 and TMUB1.

## Results

### The ERLIN complex mediates the interaction between TMUB1 and RNF170

To shed light on the processes regulated by complexes of ERLIN1 and ERLIN2, we introduced out of frame deletions in exon 1 of *ERLIN1* and exon 2 of *ERLIN2* using CRISPR/Cas9 gene editing in HeLa cells. Double knock-out (DKO) cells showed reduced mRNA levels of *ERLIN1* and *ERLIN2* consistent with RNA nonsense-mediated decay (Fig. 1A), and no detectable expression of the proteins by western blot, using antibodies that recognize regions downstream the deletions (Fig. 1B). To control for off-target effects of gene editing, we used retroviral transduction in DKO cells to re-express the proteins. For further experiments, we selected a clone that showed near to endogenous levels of ERLIN2 and substantial re-expression of ERLIN1 (DKO^+E1/2^) (Fig. 1B). Immunofluorescence using a specific antibody against ERLIN2 showed a staining consistent with localization to the ER that was lost in the DKO and restored in DKO^+E1/2^ cells (Fig. 1C). Furthermore, recovery of detergent resistant membranes (DRMs) after solubilization with a non-ionic detergent (Triton-X100) by flotation using an Optiprep gradient revealed accumulation of both endogenous and re-expressed ERLIN1 and ERLIN2 in the same fraction as other DRM markers, such as flotillin (FLOT1) and KIDINS220 (Fig. 1D). This result agrees with the known accumulation of ERLINs in cholesterol-rich lipid nanodomains (Browman et al., 2006).

**Figure 1.**
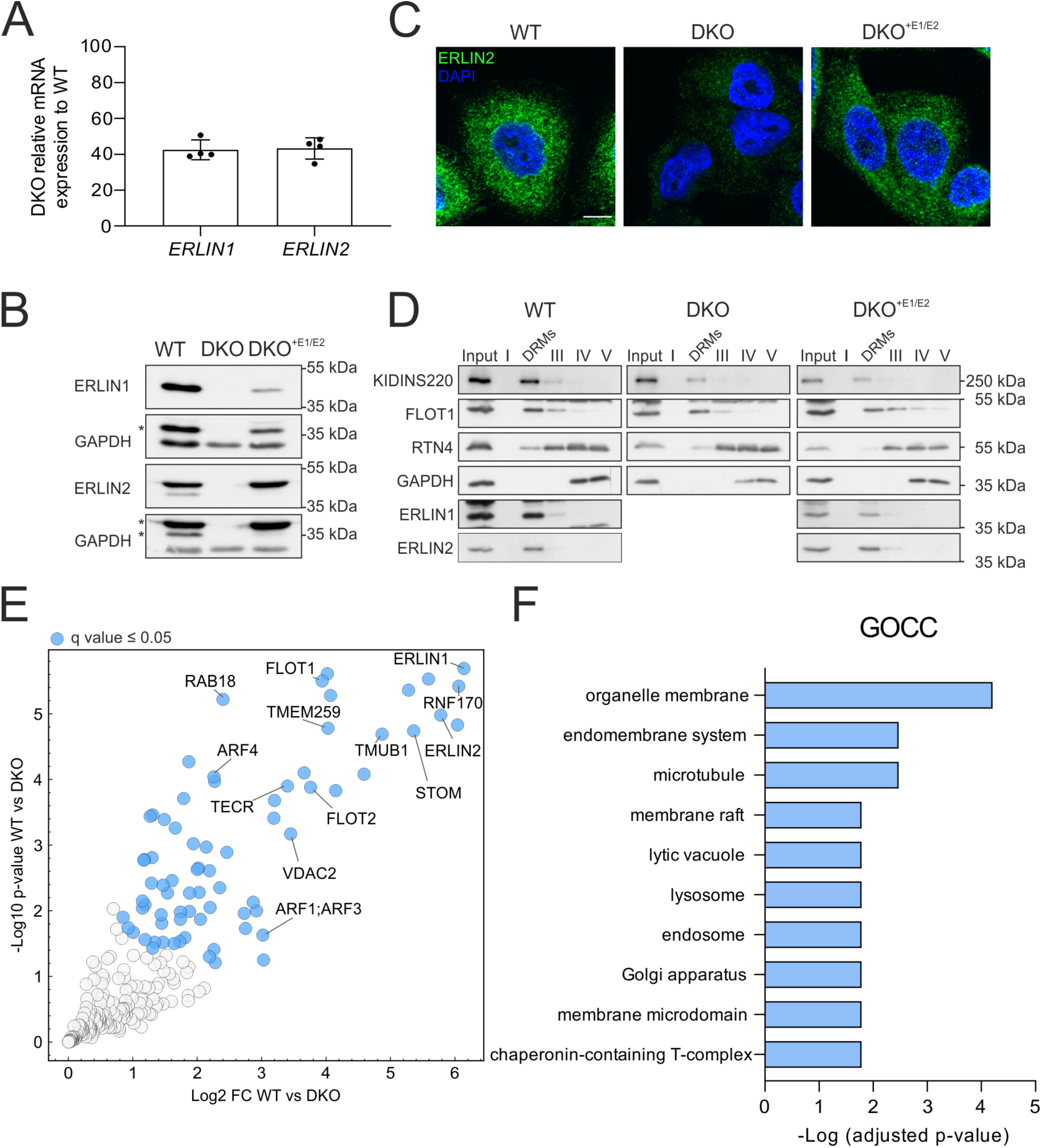
Generation of cells lacking ERLIN1 and ERLIN2 and identification of the interactome of ERLIN2. (**A**) Expression levels of *ERLIN1* and *ERLIN2* mRNAs measured by q-RT-PCR normalized for *GAPDH* in DKO compared to WT (set to 100). **(B)** Western blot for ERLIN1 and ERLIN2 in WT, DKO and DKO^+E1/E2^. GAPDH was used as loading control. Asterisks mark unspecific bands. **(C)** Immunofluorescent staining of ERLIN2 in WT, DKO and DKO^+E1/E2^. **(D)** Western blot for ERLIN1 and ERLIN2 in the input and different fractions of DRM isolating gradients from WT, DKO and DKO^+E1/E2^ cells. FLOT1 and KIDINS220 are DRM markers, RTN4 is a non-DRM membrane protein and GAPDH is a soluble protein. **(E)** Volcano plot of endogenous ERLIN2 interactors in WT compared to DKO cells. N= 4 biological replicates. Significantly enriched proteins (q value ≤0.05) in DKO are labelled in blue. **(F)** Enriched GO cellular component (GOCC) terms of proteins with q value ≤0.05 performed using the gProfiler webtool, all proteins identified in the analysis were used as background.

Currently, it is unclear if different ERLIN complexes exist in association with distinct E3 ubiquitin ligases or if ERLINs organize membrane domains containing more than one E3 ubiquitin ligase. A limitation of earlier studies is that they have been performed in overexpression conditions, which may promote unspecific interactions. We employed anti-ERLIN2 antibodies to pull down the endogenous complex and identify interacting proteins by label-free quantitative mass spectrometry (MS) (Fig. 1E, Supplementary Table 1). Among the most enriched proteins in the immunoprecipitation (IP) of WT cells were the molecular partner ERLIN1 and the E3 ubiquitin ligase RNF170. In addition, two other ERAD regulators, TMEM259 (also known as membralin), and TMUB1 were highly and significantly enriched, in agreement with previous studies (Fig. 1E) (Jo et al., 2011b; van de Weijer et al., 2020). We also identified the trans-2,3-enoyl-CoA reductase (TECR), an endoplasmic reticulum multi-pass protein that is involved in the fatty acid elongation cycle. In addition, ERLIN2 IP included other proteins that associate with DRMs, such as flotillin-1 (FLOT1) and stomatin (STOM), or proteins that traffic along the lipid rafts, as revealed by pathway analysis (Fig. 1F). We also detected proteins belonging to other organelles, such as the mitochondrial VDAC2, the peroxisomal ABCD3, and the small GTPases ARF1 and ARF4, which localize at the Golgi and are involved in vesicular trafficking (Adarska et al., 2021), possibly reflecting the presence of ERLINs at the contact sites between the ER and other organelles.

TMUB1 is a ubiquitin-like domain containing protein, which acts as an escortase stabilizing intermediates of membrane proteins during retro-translocation and delivering them to p97/VCP (Wang et al., 2022). A long and a short isoform of TMUB1 (from now on TMUB1-L and TMUB1-S) deriving from alternative initiation of translation have been previously reported (Fig. 2A) (Castelli et al., 2014; van de Weijer et al., 2020; Wang et al., 2022). Reciprocal co-immunoprecipitation experiments demonstrated that ERLIN2 interacts specifically with TMUB1-L (Fig. 2B, C). Furthermore, the amount of TMUB1-L recovered in the DRM fraction was decreased in DKO cells and rescued by re-expression of the ERLINs (Fig. 2D-F), suggesting a role of ERLIN complexes to cluster TMUB1-L in specific membrane nanodomains. In contrast, the distribution of TMUB1-S and of other ERLIN2 interactors remained unaffected (Fig. 2D-F and Fig. S1). Unfortunately, we could not test if lack of ERLINs impairs targeting of RNF170 to the DRMs because of lack of suitable antibodies.

**Figure 2.**
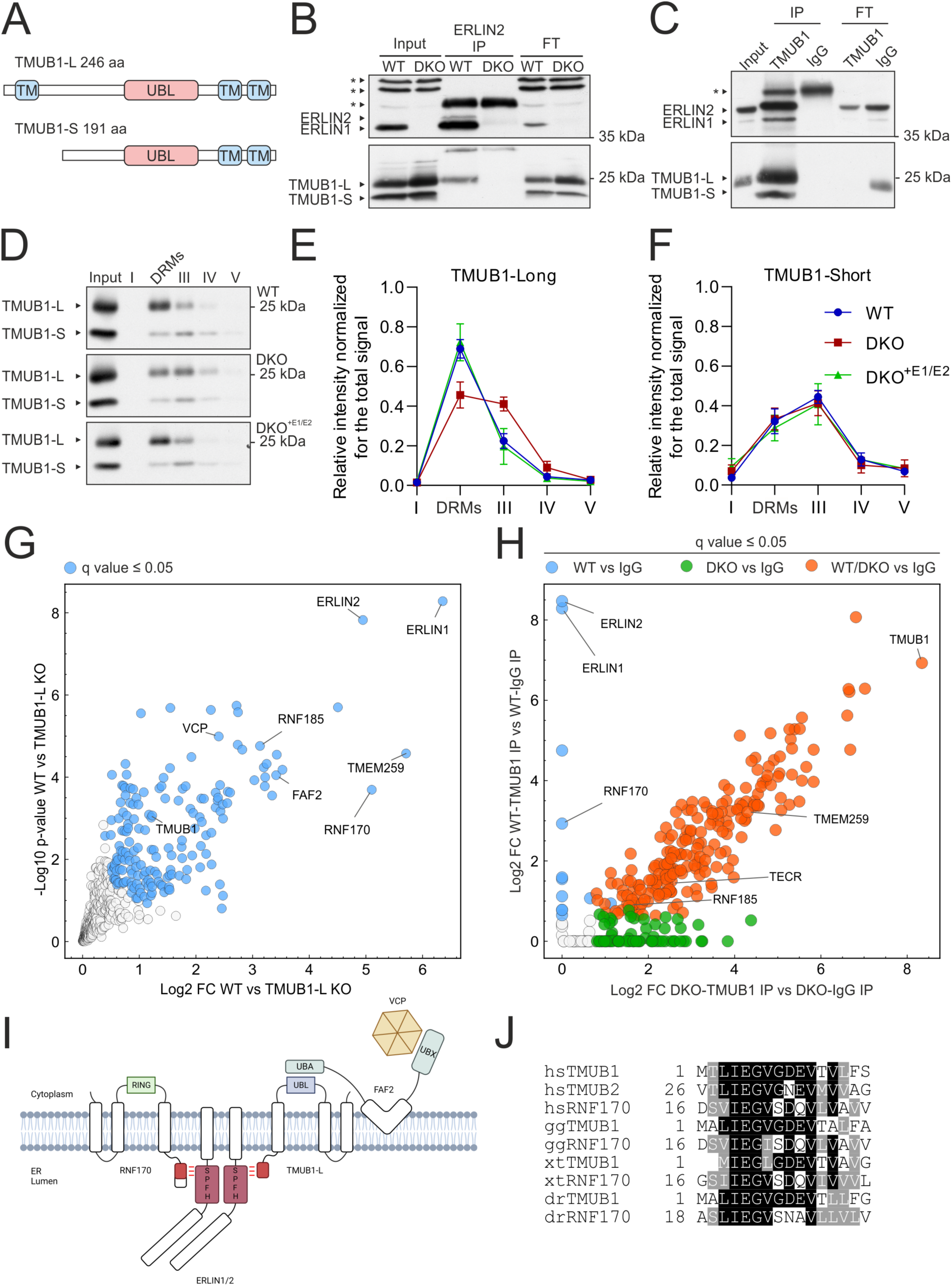
ERLINs interact with TMUB1-L and mediate the interaction of TMUB1-L with RNF170. (**A**) Scheme of the domain structure of the two human TMUB1 isoforms. **(B, C)** Western blots of reciprocal co-IPs of endogenous ERLIN2 and TMUB1 in HeLa cells. Input is equal to 10% of the proteins used for the IP. Asterisks mark unspecific bands. **(D)** Western blot of TMUB1 in the input and different fractions of DRM isolating gradients in WT, DKO and DKO^+E1/E2^. **(E, F)** Quantification of TMUB1-L (E) and TMUB1-S (F) in the different fractions. N= 4 biological replicates. **(G)** Volcano plot of endogenous TMUB1 interactors in WT compared to TMUB1-L KO cells. N= 4 biological replicates. Significantly enriched proteins (q value ≤0.05) in WT are labelled in blue. **(H)** Correlation of protein fold changes of a TMUB1 IP in WT versus ERLINs DKO cells. Significantly enriched proteins (q value ≤0.05) only in WT cells are labelled with blue, only in DKO with green, and in both conditions with orange. **(I)** Schematic representation of the ERLINs-RNF170-TMUB1-L-FAF2-VCP complex topology. **(J)** N-terminal conserved motif alignment in TMUB1-L, TMUB2 and RNF170.

To gain further insights into the specific function of TMUB1-L, we generated a TMUB1-L specific KO cell line using an inactive Cas9 fused with an adenine deaminase to mutagenize the first ATG (Gaudelli et al., 2017) (Fig. S2A), and used it a control in immunoprecipitation experiments, followed by MS (Supplementary Table 1). ERLIN1 and ERLIN2 emerged as the top proteins co-immunoprecipitated by TMUB1-L, validating our earlier findings (Fig. 2G). Moreover, we confirmed the interaction of TMUB1-L with RNF170, TMEM259, RNF185, FAF2 and VCP (Wang et al., 2022). FAF2 is anchored to the ER by a hairpin domain, contains both an UBA and UBX domain, and has been previously implicated in ERAD pathways (Mueller et al., 2008; Olzmann et al., 2013). Intriguingly, AlphaFold Multimer predicts not only the known interaction of the UBX domain with VCP (Schuberth and Buchberger, 2008), but also a possible interaction between the UBA domain of FAF2 and the UBL of TMUB1-L (Fig. S2B, C).

We then asked whether any of the molecular interactions of TMUB1-L depends on the presence of ERLIN scaffolds. To this purpose, we used TMUB1 antibodies (detecting both TMUB1 isoforms) for IP experiments in WT and DKO cells (Fig. 2H, Fig. S2D, and Supplementary Table 1). Many proteins enriched in TMUB1 IP belonged to a nucleolar compartment but were likely due to a cross-reactivity of the antibody, since a nucleolar staining could not be suppressed in cells downregulated by siRNA against TMUB1 (Fig. S2E, F). Surprisingly, lack of ERLINs completely prevented the interaction of TMUB1 with RNF170 (Fig. 2H), while the TMUB1-RNF185-TMEM259 interaction previously reported could still form in DKO cells. We conclude that the ERLIN scaffolds are required to cluster TMUB1-L and RNF170 in ER membrane nanodomains.

### A conserved luminal domain in TMUB1-L and RNF170 interacts with the ERLIN complex

TMUB1-L and RNF170 have a similar topology with a luminal domain, three transmembrane domains and a cytosolic UBL and RING domain, respectively (Fig. 2I). Inspection of the N-terminal luminal domain of TMUB1-L, which is required for interaction with ERLINs, revealed the presence of a stretch of seven amino acids that was conserved in the N-terminal domain of RNF170 (Fig. 2J). We used AlphaFold Multimer to model the three-dimensional interaction between TMUB1-L, RNF170, and a dimer of ERLIN1 and ERLIN2 (Fig. 3A). The best model predicted that the conserved N-terminal motif of RNF170 and TMUB1 interacts with the SPFH domain of each ERLIN subunit at different interfaces. The first interaction occurs with a β sheet in the ERLINs. Hydrogen bonds are predicted between Tyr51 of ERLIN1 and Gly21 of RNF170, while Phe49 and Leu51 in ERLIN2 contact Gly6 and Gly10 of TMUB1, respectively. On the second interface, Glu5 of TMUB1-L or the corresponding Glu20 of RNF170 binds to the adjacent ERLIN subunit with hydrogen bonds predicted with Arg38 and Thr44 in ERLIN1 and Arg 36 and Thr 42 in ERLIN2.

**Figure 3.**
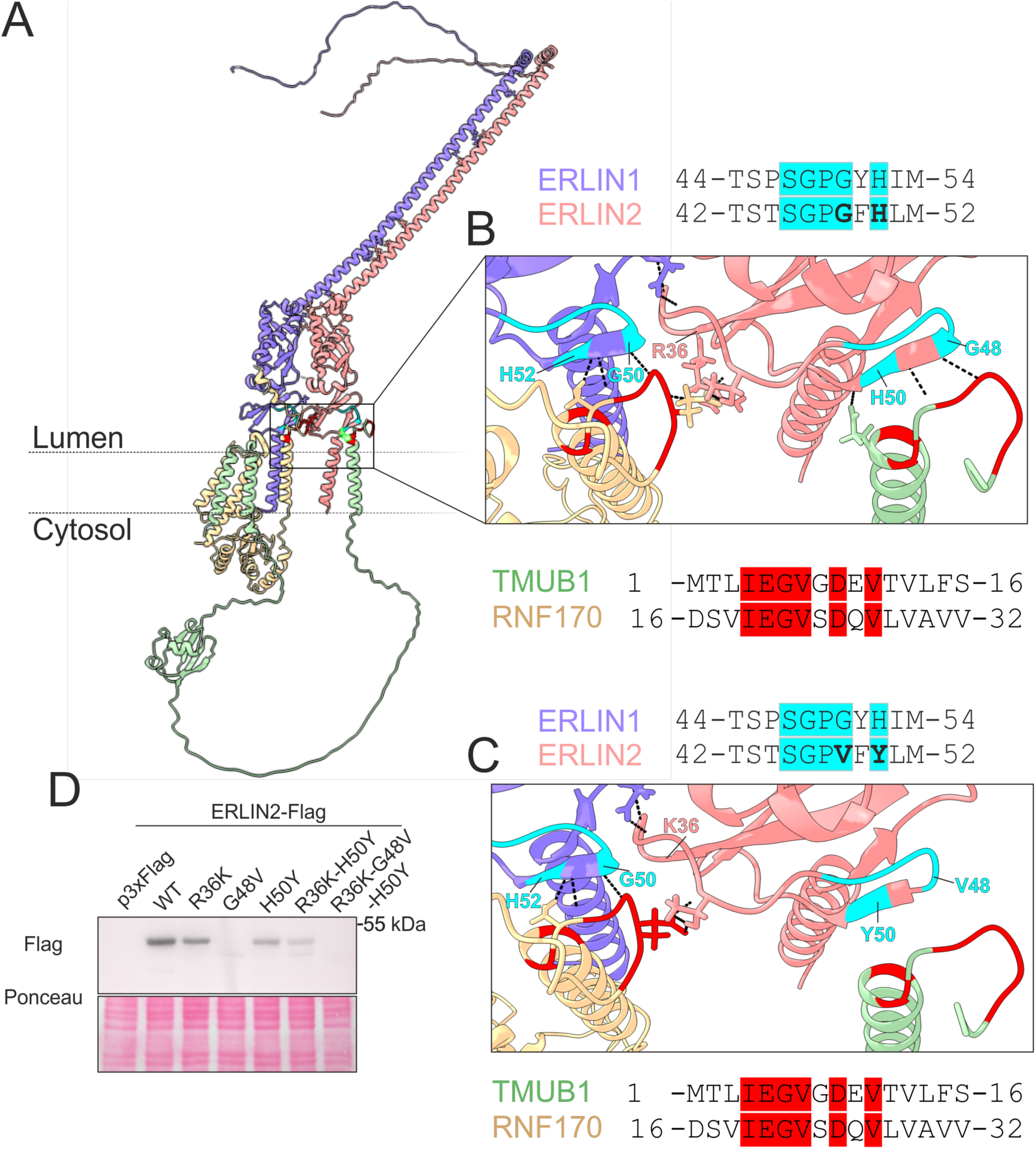
Model of the ERLINs-RNF170-TMUB1-L complex. (**A**) AlphaFold Multimer model of ERLINs-TMUB1-L-RNF170. ERLIN1 is represented in violet, ERLIN2 in pink, TMUB1 in green and RNF170 in yellow. **(B)** Enlargement of the docking site with predicted H-bonds. Conserved sequences between ERLIN1 and 2 are shown in cyan and between TMUB1 and RNF170 in red. **(C)** Enlargement of the same complex containing the ERLIN2 R36K-G48V-H50Y variants. **(D)** Western blot of transfected ERLIN2-Flag variants expressed in HeLa cells detected with anti-Flag antibody.

A Gly50Val substitution in *ERLIN1* has been implicated in autosomal recessive SPG62 (Novarino et al., 2014), while His50Tyr (ClinVar, ID: 947803) and Arg36Lys (ClinVar, ID: 1720746) in *ERLIN2* have been reported as variant of unknown significance in individuals with spastic paraplegia. We explored if these variants alter the interaction of the ERLINs with TMUB1-L and RNF170, by modelling the interaction after mutating the corresponding residues in ERLIN2 (Fig. 3C). The size of the ERLIN2 β-sheet between Gly48 and His50 is shortened by introducing the mutations, and the hydrogen bonds predicted between TMUB1-L and the ERLIN subunits at the two interaction interfaces are lost (Fig. 3C). Expression of ERLIN2-FLAG mutant constructs carrying these variants individually or in combination showed that the Gly48 is required for the stability of the protein (Fig. 3D). Similarly, the simultaneous mutation of the two residues predicted to form hydrogen bonds with both TMUB1-L and RNF170 also induces protein degradation (Fig. 3D). These data support the hypothesis that these variants are pathogenic and that the interaction of ERLINs with TMUB1-L and RNF170 plays a role in stabilizing the ERLIN complex.

### Loss of ERLINs perturbs pathways linked to cell adhesion and vesicular trafficking

To uncover pathways regulated by the ERLIN complex, we first used unbiased omics approaches. Transcriptomics analysis of WT and DKO cells confirmed the literature data showing the activation of the SREBP pathway upon ERLINs depletion (Huber et al., 2013), however this activation was milder than previously reported (Fig. 4A and Supplementary Table 2). Over-representation analysis (ORA) of DKO transcriptomes showed a significant downregulation of GO terms related to the ER, but also to focal adhesion, cell adhesion, and cadherin binding (Fig. 4B). We used label-free quantitative mass-spectrometry (MS) to reveal changes at the proteome level in DKO cells. We only considered proteins measured by at least 2 peptides, showing significant changes in abundance (ANOVA <0.05) and rescued by re-expressing the ERLINs. Out of 2,742 proteins measured in the post-nuclear supernatant, only 34 fulfilled these criteria (Fig. 4C; Supplementary Table 1). Many of these proteins functionally belonged to pathways related to sterol and lipid metabolism and cell adhesion. Most of these changes were mirrored by transcriptional alterations. ER-resident enzymes involved in neutral lipid and cholesterol metabolism (ACSL1, HMGCR and SQLE) and regulated by SREBPs were increased. Furthermore, we found that β-catenin (CTNNB1) was increased in DKO cells and rescued in the DKO^+E1/2^ cells. Alteration of the β-catenin pathway was further confirmed via western blot (Fig. S3A, B), and was paralleled by a migration defect in a scratch assay, in which DKO cells took more time to fill a gap on a plate compared to WT and DKO^+E1/2^ cells (Fig. S3C, D).

**Figure 4.**
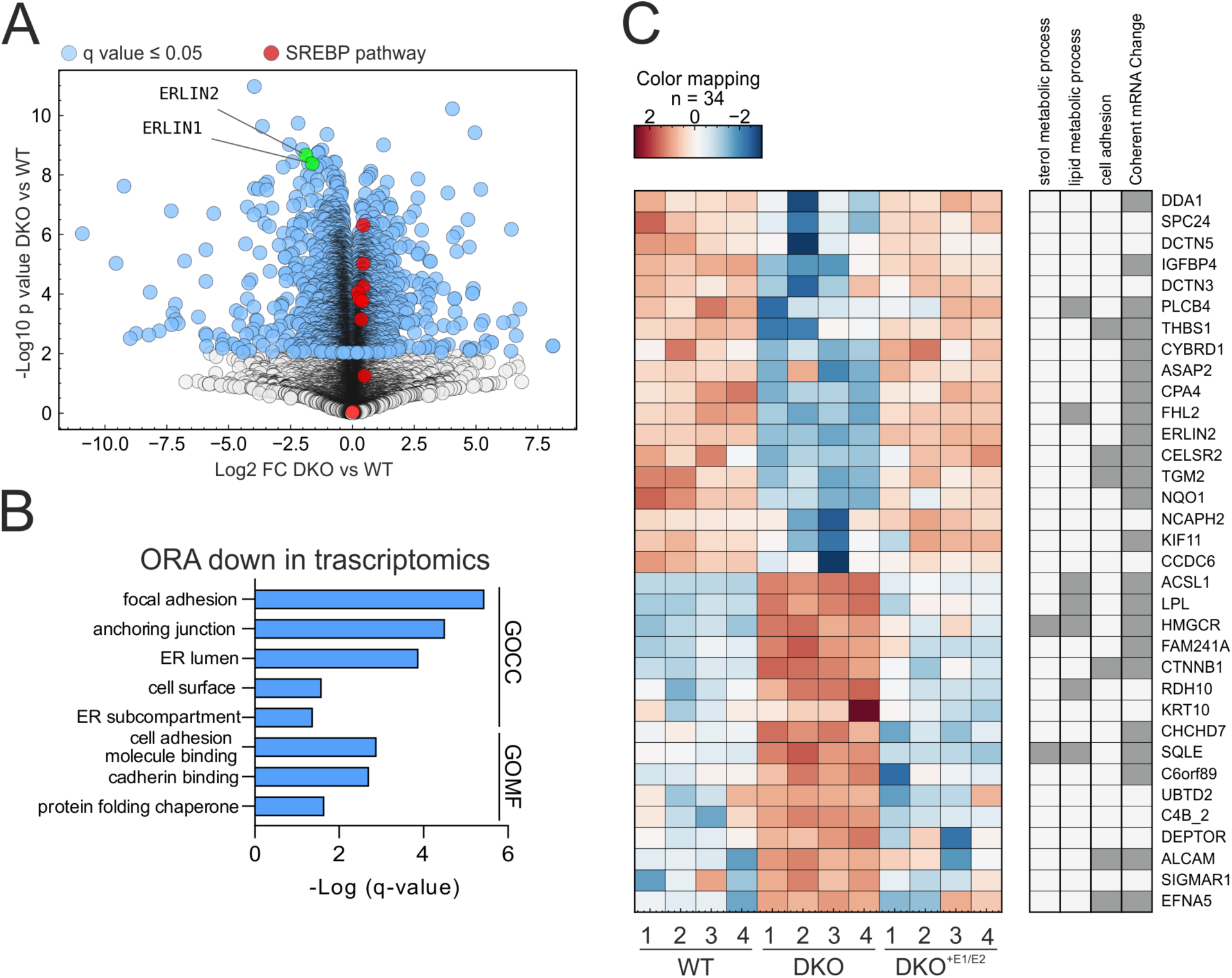
RNAseq and proteomics of DKO reveal perturbed pathways. (**A**) Volcano plot of the RNA-seq analysis. Genes with q value ≤0.05 are labelled in blue and known SREBPs target genes are labelled in red. **(B)** Over-representation analysis (ORA) of the downregulated genes from the RNA-seq. N= 4 biological replicates. **(C)** Heatmap of z-score normalized intensities of significant proteins in ANOVA (q value ≤0.05) with absolute fold changes > 1.5 in WT vs DKO and DKO vs DKO^+E1/E2^ but not changed in WT vs DKO^+E1/E2^. N= 4 biological replicates.

We reasoned that the lack of the scaffolding function of the ERLINs complex in the ER may not necessarily affect overall protein abundance but may change their dynamic subcellular distribution by impairing recruitment to cholesterol-enriched nanodomains. We therefore combined the flotation gradient used to separate DRMs with stable isotope labelling by amino acids in cell culture (SILAC) (Ong et al., 2002) and analysed protein abundance in five collected fractions with different densities. The major advantage of this approach is allowing us to combine WT, DKO and DKO^+E1/2^ cell lysates before loading them on the flotation column, thus reducing the variability of fraction collection among samples (Fig. 5A). Proteins associated with DRMs are recovered in low-density fractions, however they can shift from one fraction to another upon altered cellular lipid composition or signalling stimulation (Foster et al., 2003; Simons and Toomre, 2000). This approach confirmed that in WT cells, ERLIN1, ERLIN2, and other DRM-associated proteins, such as FLOT1 and FLOT2, are enriched in fraction 2, as previously detected by western blot (Fig. S4 and Fig. 1D). The overall distribution of the log2 normalized protein ratio was similar between all comparisons in fractions II to V. Fraction I contained significantly less proteins than the other fractions, which accumulated in the DKO compared to the other genotypes. We used unsupervised clustering to identify proteins that were significantly changed in the three comparisons (KO/WT; DKO^+E1/2/DKO^, and DKO^+E1/2/^WT). We identified Cluster 170 in fraction I and cluster 155 in fraction II as containing proteins that were upregulated in the DKO and rescued in the DKO^+E1/2^ cells (Fig. 5C, D, F, G, and supplementary Table 3). Pathway analysis showed that these clusters were enriched in proteins belonging to adherens and anchoring junctions, to the plasma membrane, and to membrane-bound vesicles (Fig. 5E, H). Surprisingly, cluster 577 in fraction IV contained proteins belonging to the same GO terms, showing however an opposite trend compared to the previous two clusters. Thus, proteomics suggests perturbations of proteins that follow the secretory route.

**Figure 5.**
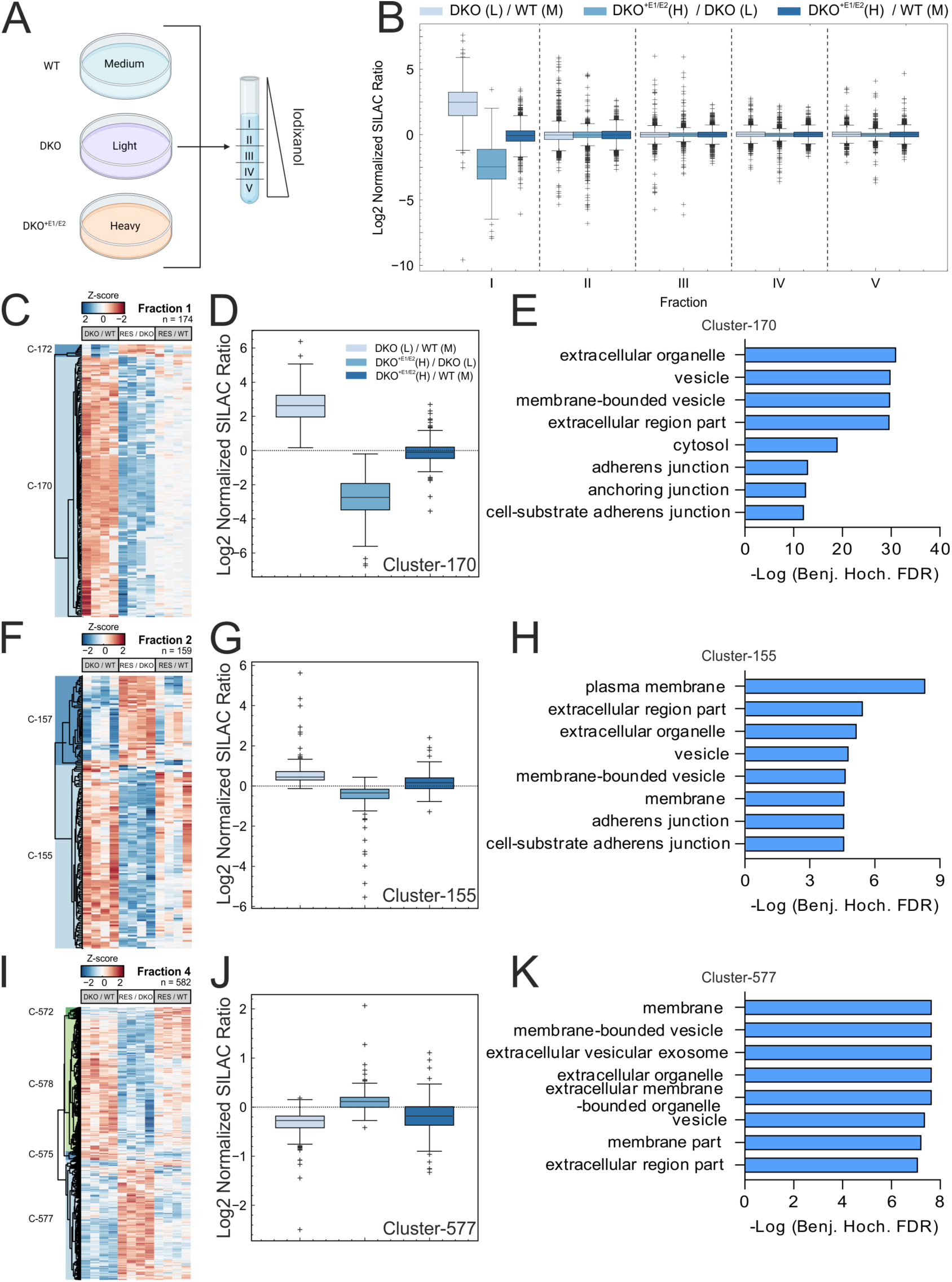
Lack of ERLINs alters the distribution of proteins in DRM fractions. (**A**) Scheme of the SILAC approach combined with the DRMs isolation protocol. **(B)** SILAC normalized ratio of each fraction. **(C-K)** Unsupervised cluster analysis of significantly regulated proteins (ANOVA q-value σ 0.05) detected in fractions 1, 2 and 4 reveals clusters of proteins that are perturbed in DKO and rescued in DKO^+E1/E2^ (RES). Heat maps of fractions 1, 2 and 4, respectively (C, F, I). Log2 normalized SILAC ratio of clusters 170, 155, and 577 (D, G, J) and corresponding significantly enriched GO terms (q-value σ 0.05) with the Fisher’s Exact test (E, H, K).

### Loss of the ERLIN complex leads to a collapse of ER tubules and Golgi fragmentation

Based on the previous results, we hypothesized that trafficking along the secretory compartment may be impaired in DKO cells. As a first step, we used antibodies against reticulon 4b (RTN4b), which is enriched in the tubular ER, to assess if the general morphology of the ER was affected in DKO cells. Intriguingly, DKO cells were characteristically depleted of peripheral tubular ER in comparison to both WT and DKO^E1/2^ cells (Fig. 6A, B). We noticed that the cell area labelled by RTN4b staining in DKO cells appeared smaller 24 hours after plating (Fig. S5A), despite the cell size was unchanged when assessed in cytofluorimetry (Fig. S5B). Therefore, the ER phenotype may be influenced by a defect of the cells in spreading on the plate, consistent with the cell adhesion defect.

**Figure 6.**
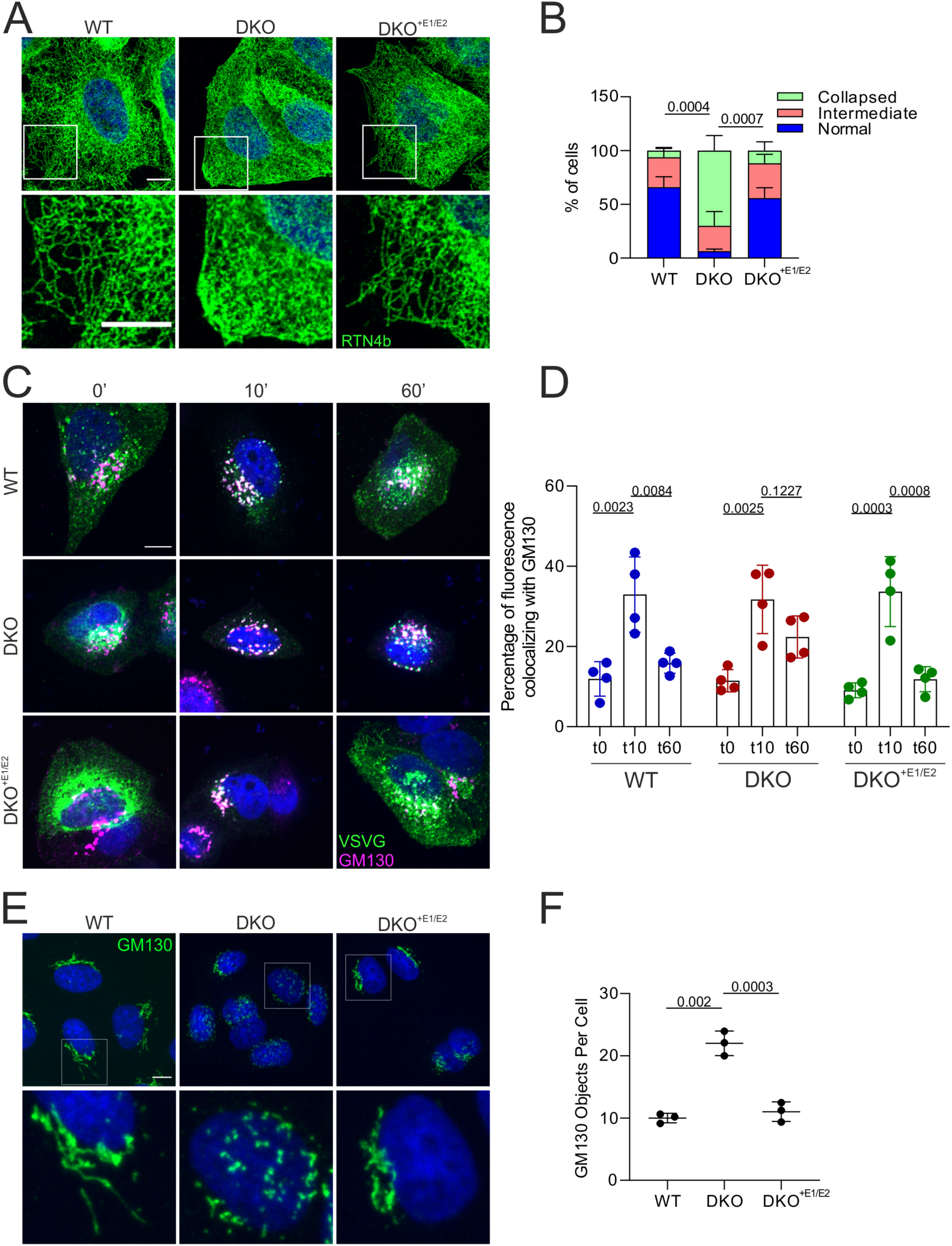
DKO cells show defects along the secretory pathway. (**A**) Immunofluorescence of the ER using an anti-RTN4 antibody in cells of different genotypes. **(B)** For quantification of A, ER tubules were classified as normal, intermediate and collapsed. N= 3 independent experiments (at least 90 cells were analysed per genotype). **(C)** RUSH assay using the VSVG-GFP construct. Cells were fixed at indicated time points and stained for GM130 to label the Golgi apparatus. **(D)** Quantification of the percentage of GFP colocalising with GM130 at each time point. The ratio between the GFP signal inside the GM130 positive area and the total GFP signal is shown. N= 4 biological replicates (at least 110 cells were analysed per genotype for each time point). **(E)** Immunofluorescence of the Golgi apparatus using an anti-GM130 antibody. **(F)** Quantification of experiment as in E. N= 3 biological replicates (at least 300 cells were analysed per genotype) and the statistical test shown is the one-way analysis of variance with post Tukey’s multiple comparison test.

To assess trafficking along the secretory system, we employed the retention using selective hooks (RUSH) system (Boncompain et al., 2012). This system comprises a hook fused to streptavidin, which is bound to the donor compartment (the ER) and a reporter protein fused to a streptavidin binding domain (SBD). Addition of biotin to the medium synchronously competes with the SBD, releasing the reporter protein from the hook and allowing its trafficking along the secretory pathway. We employed the vesicular stomatitis virus G (VSVG) glycoprotein expressed with a fluorescent EGFP tag as a reporter and monitored in live imaging its trafficking after biotin addition in WT, DKO and DKO^+E1/2^ cells (Fig. S5C and Supplementary videos 1-3). In WT and DKO^+E1/2^ cells, VSVG-GFP translocated from the ER to the Golgi compartment within approximately 10 min, followed by trafficking to the exocytic compartment. In ERLIN-deficient cells, VSVG-GFP moved from the ER to a fragmented vesicular compartment within 10 min after adding biotin, and then remained largely confined within these structures. When we analysed co-localisation of the VSVG signal with a Golgi marker in cells fixed at different time-points after biotin addition, we confirmed that in DKO cells the VSVG-EGFP remained elevated in the Golgi after 60 min from biotin addition, in contrast to WT and DKO^+E1/2^ cells (Fig. 6C, D). Consistently, loss of ERLINs induced Golgi fragmentation (Fig. 6E, F), a phenotype that was completely rescued in DKO^+E1/2^ cells (Fig. 6E, F). Dispersion of the Golgi apparatus was also observed upon TMUB1-L depletion (Fig. S5D, E). Thus, ER-to-Golgi trafficking appeared unaffected by the lack of the ERLIN complex, while the Golgi apparatus showed a morphology defect and post-Golgi trafficking was impaired.

### Excessive cholesterol esterification in absence of ERLINs causes Golgi fragmentation

The Golgi apparatus is a central sorting station not only for proteins, but also for lipids that traffic along the secretory pathway. While the presence of lipid rafts in the ER is still debated, it is accepted that lipid nanodomains enriched with cholesterol and sphingolipids are present along the secretory pathway, with increasing abundance from the ER to the plasma membrane. These domains are implicated in the intracellular trafficking of proteins, such as GPI-linked receptors, integrins, and other adhesion molecules. By binding cholesterol at the ER, complexes of ERLINs may control the distribution of cholesterol in cellular membranes. We measured levels of ceramides and sphingomyelin in the DRMs of WT, DKO, and DKO^+E1/2^ cells using targeted lipidomics. We found higher quantity of species of both lipids containing fatty acids with chain lengths between C18 and C24 in the DKO cells, which were restored to WT levels in the DKO^+E1/2^ cells (Fig. 7A, B and supplementary Table 4). The protocol used for the preparation of DRMs prevented measuring cholesterol levels in these fractions. We therefore assessed the content of free cholesterol and cholesterol esters (CE) in the total cell lysates. Unesterified cholesterol was not affected upon ERLINs depletion (Fig 7C). However, total CE and especially the abundant endogenously synthesized CE 18:1 species were significantly increased when ERLINs were depleted and recovered to WT levels in DKO^+E1/E2^ cells (Fig. 7D, E and Supplementary Table 4). Levels of triacylglycerols (TAGs) also increased in the DKO compared to both WT and DKO^+E1/2^ cells (Fig. 7F). Interestingly, this accumulation involved mainly TAGs with long chain fatty acids (LCFAs), while was less prominent in TAGs with very long chain fatty acids (VLCFAs, carrying more than 20 C) (supplementary Table 4). Consistent with the accumulation of neutral lipids, larger lipid droplets (LDs) labelled by Bodipy 493/503 accumulated in DKO cells, a phenotype that was rescued upon ERLINs re-expression (Fig. 7G).

**Figure 7.**
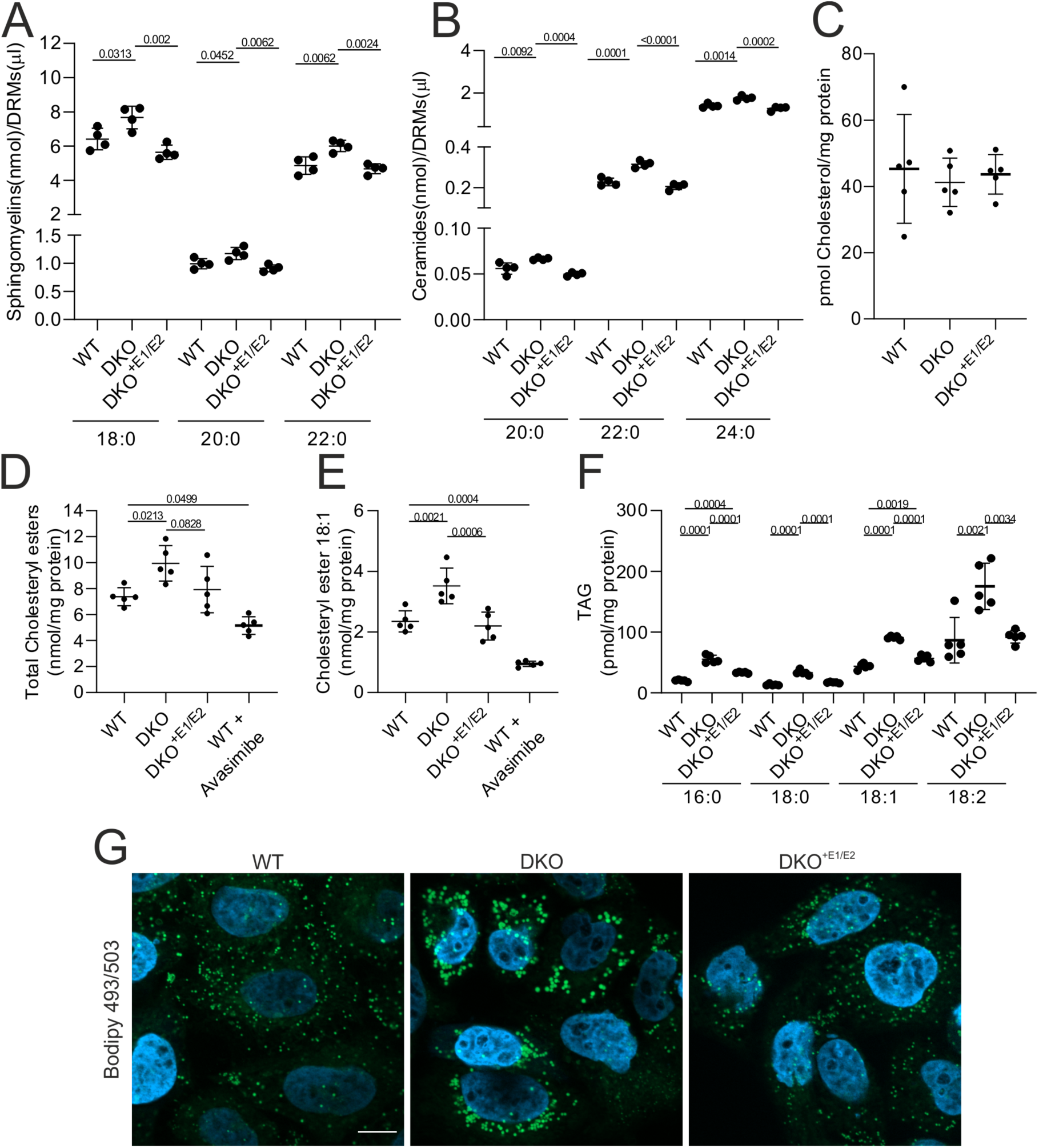
DKO cells show sphingolipid accumulation in DRMs and neutral lipid accumulation in LDs. (**A, B**) Sphingomyelins (A) and ceramides (B) accumulate in DRMs of DKO cells. N= 4 biological replicates. Only significant species are shown. **(C-F)** Lipidomics analysis of cholesterol (C), total cholesteryl ester (D), cholesteryl ester 18:1 (E) and TAGs (F). When indicated, WT cells were treated with avasimibe to confirm endogenous synthesis. N= 5 biological replicates. Statistical test shown is the one-way analysis of variance with post Tukey’s multiple comparison test. **(G)** Staining of LDs with Bodipy 493/503.

Esterification of excess cholesterol at the ER and its deposition in LDs are observed when cholesterol transport from the ER to the Golgi is impaired. Cholesterol trafficking between these two organelles mostly occurs via non-vesicular transport, at sites of close apposition between the membranes of the organelles (Ikonen and Zhou, 2021). This transfer is mediated by oxysterol-binding proteins (OSBPs), which bind cholesterol at the ER and release it at the Golgi in exchange for a molecule of phosphatidylinositol-4-phosphate (Ikonen and Zhou, 2021). The FFAT domain of OSBP binds VAPA at the ER, while the PH domain binds Arf1 at the Golgi membrane. Notably, Arf1 was identified in the ERLIN2 interactome (Fig. 1E), suggesting close apposition between the ERLIN complex and the Golgi. Cholesterol balance in the Golgi is essential for protein transport, and both cholesterol depletion and cholesterol loading block trafficking of secretory proteins from the trans-Golgi (Stüven et al., 2003). In addition, lack of cholesterol in the Golgi may lead to vesiculation (Stüven et al., 2003). DKO cells showed a modest but significant SREBPs activation, indicating that they evade feedback inhibition by cholesterol synthesis. While the destabilization of INSIG1 upon ERLINs knockdown could explain this result (Huber et al., 2013), an alternative explanation could be that increased cholesterol esterification maintains low levels of cholesterol in the ER, impairs transport to the Golgi, and prevents feedback inhibition of SREBPs. To test this hypothesis, we treated WT and DKO cells with the SOAT1 inhibitor avasimibe. Remarkably, inhibition of SOAT1 activity not only reduced the size of the LDs in DKO cells (Fig. 8A, C), but also rescued the Golgi fragmentation (Fig. 8B, D), supporting our hypothesis. SOAT1 inhibition was sufficient to revert the transcriptional upregulation in DKO cells of 3-hydroxy-3-methylglutaryl-CoA synthase (HMGCS), a SREBP target gene (Fig. 8E). We propose a model in which the ERLIN complex, by binding cholesterol, restricts its esterification, and thus promotes transport to the Golgi. Loss of ERLIN1/2 scaffolds on the one side depletes the ER of cholesterol, activating the SREBP pathway, and on the other side leads to Golgi vesiculation and defects in post-Golgi trafficking.

**Figure 8.**
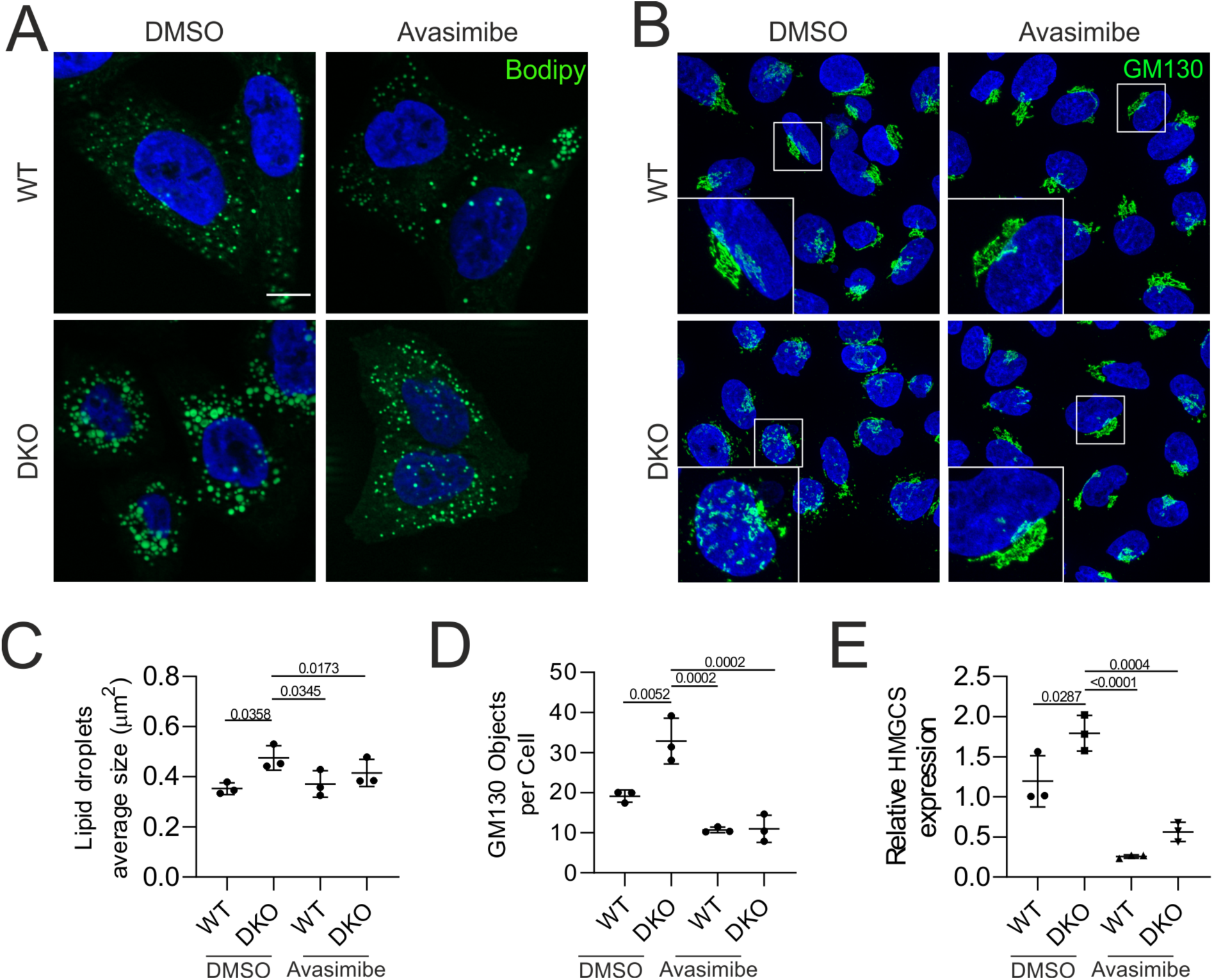
Avasimibe treatment of DKO cells rescues LD accumulation, Golgi fragmentation, and SREBP activation. (**A**) Bodipy 493/503 staining and **(B)** GM130 immunofluorescence of WT and DKO cells treated with and without Avasimibe. **(C)** Quantification of LD size. N= 3 biological replicates (at least 130 cells were analysed per genotype). **(D)** Quantification of Golgi fragmentation. N= 3 biological replicates (at least 340 cells were analysed per genotype). **(E)** Relative *HMGCS* expression in WT and DKO cells with and without Avasimibe. GAPDH was used as control. The statistical test shown is the one-way analysis of variance with post Tukey’s multiple comparison test.

## Discussion

Our study identifies a key role of ERLIN1 and ERLIN2 scaffolds on the ER to mediate the interaction between TMUB1-L and RNF170 and to limit cholesterol esterification, thereby facilitating transport of cholesterol to Golgi. Lack of the ERLIN complex impairs the secretory pathway, leading to the collapse of the tubular ER, Golgi fragmentation, impaired post-Golgi trafficking, and cell migration defects. These results suggest novel pathogenic pathways underlying the degeneration of long axons of the corticospinal tracts in patients carrying mutations in *ERLIN1* or *ERLIN2*.

We show that the ERLIN complex functions as a platform to recruit the E3-ubiquitin ligase RNF170 and the long isoform of TMUB1. This interaction occurs via a conserved eight amino acid motif in the N-terminus of RNF170 and TMUB1-L, which can bind adjacent ERLIN subunits in a hetero-oligomeric complex, based on three-dimensional modelling using AlphaFold Multimer. Consistently, lack of the N-terminal motif in TMUB1-S abrogates binding to the ERLINs, TMUB1 immunoprecipitates RNF170 in WT but not DKO background, and less TMUB1-L is recovered in the DRM fraction in the DKO. The motif is unique for RNF170 and TMUB1 and is conserved in TMUB2, but not in others E3 ubiquitin ligases previously linked to ERLINs. The interaction interface on the ERLIN side involves two luminal regions at the beginning of the SPFH domain. Particularly relevant is a β sheet formed by a glycine, followed by an aromatic amino acid (Tyr in ERLIN1 and Phe in ERLIN2), and a histidine. Shortening of this β sheet in ERLIN2 by mutating the glycine to valine to reproduce a HSP mutation found in *ERLIN1* (Novarino et al., 2014) results not only in the loss of the predicted hydrogen bond with TMUB1 and RNF170, but also in the destabilization of the protein, without impairing the protein oligomerization. A similar effect was obtained when mutating at the same time the two ERLIN2 binding interfaces, by introducing mutations reported as variant of unknow significance in patients with HSP. Based on the AlphaFold modelling and the experimental results, our data support the pathogenicity of these mutations, and substantiate the relevance of this interaction.

Our data are consistent with a core complex composed of ERLINs, RNF170 and TMUB1-L. Modelling predicts that both TMUB1-L and RNF170 bind the ERLIN scaffold at the same time, however we cannot exclude the presence of different complexes containing one or the other protein. In contrast to RNF170, TMUB1 still binds RNF185 and TMEM259 in absence of ERLINs. A previous study showed that the interaction of TMUB1 with RNF185 is not direct but mediated by TMEM259, and that ERLINs were not essential for ERAD of substrates regulated by RNF185 (van de Weijer et al., 2020), and thus agrees with our findings. In agreement with a recent publication (Wang et al., 2022), we confirmed the interaction of TMUB1-L with VCP, and we propose that it occurs via FAF2 (Fig. 2G and Fig. S2B, C). FAF2 was also previously shown to interact with ERLIN2 upon overexpression (Wolf et al., 2021), however our data strongly argue against a direct interaction.

Oligomers of ERLINs form cup-shaped complexes that project into the lumen of the ER. It has been estimated that each ERLIN subunit can bind up to four molecules of cholesterol (Huber et al., 2013). Cells depleted of ERLINs show activation of SREBPs (Huber et al., 2013), and we confirmed this finding. It was proposed that ERLINs can directly bind INSIG1 (Huber et al., 2013), a protein that retains SCAP in the ER upon cholesterol sufficiency, thereby suppressing the trafficking of SREBPs to the Golgi, where they are proteolytically processed to release an active transcription factor that upregulates genes involved in cholesterol biosynthesis (Shimano and Sato, 2017). In this scenario, ERLINs would act as scaffolds where not only cholesterol is concentrated, but also where the INSIG1-SCAP interaction is stabilised. Our data provide an alternative explanation for how ERLIN scaffolds regulate cholesterol homeostasis. We propose that cholesterol bound to ERLINs is less accessible to SOAT1, preventing its esterification and facilitating cholesterol transport to the Golgi. Esterification of cholesterol and deposition in LDs together with triacylglycerols is a strategy by which cells detoxifies excess cholesterol, however in DKO cells unhinged cholesterol esterification would lead to cholesterol depletion on the ER, triggering SCAP-mediated movement of SREBPs to the Golgi. In support of this model, we show that inhibition of SOAT1 with avasimibe rescues both the accumulation of large LDs and the upregulation of the SREPB target gene HMGCS1 in DKO cells. Furthermore, avasimibe treatment restores the morphology of the Golgi apparatus in DKO, indicating that the Golgi vesiculation is most likely the result of cholesterol depletion in the Golgi.

Other data render our model plausible. First, cholesterol binding sites have been mapped to residues located on the luminal side of the ERLIN cups (Hulce et al., 2013), so that cholesterol may remain buried within the cavity of the complex and inaccessible to SOAT1. Second, TMUB1 was also found to bind cholesterol (Hulce et al., 2013), and SOAT1 was present in the TMUB1 interactome, supporting the proximity of this enzyme to the complex. Notably, DKO cells showed a tendency to increased levels of SOAT1 (log2FC =0.40; q =0.07), despite unchanged transcript levels. Third, the presence of Arf1 and Arf4 in the ERLIN2 IP supports the location of the ERLIN scaffolds to ER-Golgi contacts sites involved in cholesterol trafficking. Taken together, our results illustrate the role of SPFH domain-containing proteins in defining specific membrane nanodomains where lipid metabolism activities are regulated, and to which proteins are recruited or excluded. Recently, an inhibitory relationship between calcium release from the ER via IP3 receptors and cholesterol transport to the Golgi has been unravelled (Malek et al., 2021), rationalising the two-pronged function of ERLINs to coordinate ERAD of IP3 receptors via RNF170 (Lu et al., 2011) and cholesterol homeostasis.

Depletion of TMUB1-L isoform alone was sufficient to induce a Golgi fragmentation comparable with the depletion of the ERLIN complex. Intriguingly, TMUB1 and FAF2 are the closest orthologues to the yeast proteins Dsc3 and Ubx3, which are implicated in the so called Endosome and Golgi-Associated Degradation (EGAD) (Schmidt et al., 2019). This pathway regulates sphingolipid metabolism by actively extracting membrane proteins at the Golgi and delivering them to the proteasome. Whether a similar pathway exists in mammalian cells and involves TMUB1 is currently unknown. Several studies have linked Golgi vesiculation and trafficking of secretory proteins from the trans-Golgi network to cholesterol imbalance (Reverter et al., 2014; Runz et al., 2006). We used the model VSVG molecule to show that trafficking is halted at the Golgi, but the SILAC proteomics suggest a redistribution of proteins that move along the secretory pathway, supporting a more global defect in DKO cells. This involves several adhesion molecules, explaining the migration and cell spreading defects of DKO cells, and implicating perturbations of these pathways in the degeneration of long corticospinal axons in HSP caused by mutations in *ERLIN1* or *ERLIN2*. Another form of HSP (SPG5) is caused by mutations in *CYP7B1* (Tsaousidou et al., 2008), which encodes a cytochrome P450 7α-hydroxylase implicated in cholesterol and bile acids metabolism. As a result, patients accumulate 25-hydroxycholesterol and 27-hydroxycholesterol (Schule et al., 2010). The latter are potent inhibitors of SREBPs, and therefore can suppress cholesterol synthesis, potentially leading to cholesterol depletion in the Golgi (Radhakrishnan et al., 2007), and to a similar pathogenic cascade. In recent years causative HSP mutations have been identified in *RNF170* (Chouery et al., 2022; Wagner et al., 2019). Future studies will be needed to fully understand the link between unbalanced cholesterol metabolism, regulated ubiquitination at the ER, and HSP.

## Materials and methods

### Cell culture

HeLa cells were cultured in Dulbecco’s Modified Eagle Medium (DMEM) containing 4.5 g/L of glucose (Gibco, #11960044), 2 mM glutamine (Gibco, #25030024), 10,000 u/ml penicillin, 10,000 µg/ml streptomycin (Gibco, #15140122) and 10% FetalClone III (GE Healthcare Hyclone, #SH30109.03). For SILAC labelling, cells were grown in arginine and lysine free DMEM with 4.5 g/L glucose (Silantes; # 28001300), containing dialyzed FCS (Gibco, #26400044), penicillin-streptomycin-glutamine (Gibco, 10378016) and different isotopes of arginine and lysine. We used medium isotopes (Arg-6, Lys-4) (Silantes, # 201203902 and 211103913) for the WT, light (Arg-0, Lys-0) (Silantes, # 201003902 and 211003902) for the DKO and heavy (Arg10, Lys 8) (Silantes, # 201603902 and 211603902) for the DKO^+E1/E2^. Cells were regularly tested to be *Mycoplasma*-free.

### Knock out of ERLIN1 and 2 in HeLa cells via CRISPR/Cas9

ERLIN1 and ERLIN2 were knocked out using a CRISPR/Cas double nickase approach as described previously (Ran et al., 2013). CRISPR guideRNAs (gRNA) were designed to target exon1 of *ERLlN1* (guide sequence sense strand 5’-GCCATCTGGCTGTGTACTAC-3’, guide sequence antisense strand 5’-TGTGGATGGAGGCGTAGAGC-3’) and exon 2 of *ERLIN2* (guide sequence sense strand 5’-GTTGGGAGCAGTTGTGGCTG-3’, guide sequence antisense strand 5’-AGGATAAAGGCTCACTGATG-3’), respectively and confirmed with CRISPOR (http://crispor.tefor.net). Oligonucleotides were purchased from Sigma-Aldrich and cloned into the pX335 vector (Addgene #42335). Respective px335-guideRNA constructs were transfected into HeLa cells on five consecutive days and separated via serial dilution to obtain monoclones. Positive clones were identified by western blot analysis using anti-ERLIN1 and anti-ERLIN2 antibodies and confirmed by subcloning the affected gene loci into pcDNA3 (amplification of exon 1 of *ERLIN1* with 5’-TTAGGATCCGCGAGAGAGCGCCAAGTTTC-3’ and 5’-AATGCGGCCGCCAACATCTGGAGGGTGCT-3’ and amplification of exon 2 of *ERLIN2* with 5’-TTAGGATCCGAAAGGGGAACGTTGACTGA-3’ and 5’-AATGCGGCCGCCTTTCCCGTGGTTACTGACA-3’) followed by sequencing of 15 subclones, each.

### Generation of retroviruses and transduction

Coding regions of *ERLIN1* and *ERLIN2* (amplified with 5’-GGGGACAAGTTTGTACAAAAAAGCAGGCTAGAATGAATATGACTCAAGCCCG-3’ and 5’-GGGGACCACTTTGTACAAGAAAGCTGGGTCTCAACCTGTGCTCTCTTTGTTTT-3’; 5’-GGGGACAAGTTTGTACAAAAAAGCAGGCTACCATGGCTCAGTTGGGAGCAGTT-3’ and 5’-GGGGACCACTTTGTACAAGAAAGCTGGGTCTCAATTCTCCTTAGTGGCCGTCT-3’ respectively) were cloned into the pDONR221 and insert into the pBABE-puro vector using the gateway recombinant cloning (Morgenstern and Land, 1990). Respective constructs were transfected into Phoenix AMPHO packaging cells for 24 h. The harvested virus-containing medium was filtered through a 0.45 μm filter. For infection, HeLa cells at 70% of confluency were incubated for 24 h in virus-containing medium and 4 μg/ml Polybrene. Cells with positive genome integration were selected by incubation of infected cells in DMEM containing 5 μg/ml puromycin for 48 h. Expression of the gene of interest was confirmed by western blot analysis using ERLIN1 and ERLIN2 antibodies, respectively.

### TMUB1 long KO generation and selection

The start codon of the *TMUB1* gene was mutagenized with an adenine base editor in order to convert the ATG into GTG (Gaudelli et al., 2017). The gRNA was designed using the rgenome webtool (http://www.rgenome.net/), primers were purchased from Sigma-Aldrich and cloned into a gRNA Cloning Vector (gift from George Church (Addgene plasmid # 41824)) after digestion with AflII and through the Gibson assembly of the PCR product of the primer pair (guide sequence sense strand 5’-TTTCTTGGCTTTATATATCTTGTGGAAAGGACGAAACACCGCGCCATGACCCTGATTGAAG –3’, guide sequence antisense strand 5’-GACTAGCCTTATTTTAACTTGCTATTTCTAGCTCTAAAACCTTCAATCAGGGTCATGGCGC –3’). The gRNA containing vector was co-transfected with the pCMV_ABEmax_P2A_GFP and after 24h sorted using the INFLUX Cell-Sorter from the CECAD Flow cytometry facility. Two 96-well plates were prepared and the successful mutagenesis was tested via western blot analysis.

### Flow cytometry

Cell size was compared using a BD LSR-Fortessa Analyzer. 24h after seeding, cells were detached with trypsin, resuspended in fresh DMEM and washed twice with PBS. Pellets were resuspended in PBS and analysed. Cell debris and cell doublets were excluded from the analysis by gating. For the TMUB1-L KO sorting, an un-transfected control not expressing the GFP was used to discriminate the GFP positive events.

### RNA isolation, RNA-seq and qPCR analysis

RNA of cells was isolated with TRIzol® (Invitrogen, #15596026) reagent according to the manufacturer’s protocol (Life Technologies). Reverse transcription was performed using the Superscript First-Strand synthesis system (Thermo Fisher, #11904018). Real-time qPCR was performed in a thermocycler (Applied Biosystems) with SYBR® green Master Mix (Thermo Fisher, #4309155) and respective primer pairs (ERLIN1 sense primer 5’-GAAAGCTCACTCCCCTCTAAG-3’ and antisense primer 5’-TGTTCCCACTTAA CCCCTTG-3’; ERLIN2 sense primer 5’-ACGCTTCAAGAGGTCTACATTG-3’ and antisense primer 5’-ATTGCCTCTGGTATGTTGGG-3’; HMGCS sense primer 5’-CATTAGACCGCTGCT ATTCT-3’ and antisense primer 5’-AGCCAAAATCATTCAAGGTA-3’). GAPDH was used as reference gene in all analysis (GAPDH sense primer 5’-AATCCCATCACCATCTTCCA-3’ and antisense primer 5’-TGGACTCCACGACGTACTCA-3’) and the fold enrichment was calculated using the formula 2(-ΔΔCt). For RNA-seq analysis the RNA was submitted to the Cologne Genomic Center and prepared as followed. Libraries were prepared using the Illumina® Stranded TruSeq® RNA sample preparation kit. ERCC RNA Spike-In Mix (Thermo Fischer) was added to the samples before library preparation. Library preparation started with 1µg total RNA. After poly-A selection (using poly-T oligo-attached magnetic beads), mRNA was purified and fragmented using divalent cations under elevated temperature. The RNA fragments underwent reverse transcription using random primers. This was followed by second strand cDNA synthesis with DNA Polymerase I and RNase H. After end repair and A-tailing, indexing adapters were ligated. The products were then purified and amplified (15 PCR cycles) to create the final cDNA libraries. After library validation and quantification (Agilent Tape Station), equimolar amounts of library were pooled. The pool was quantified by using the Peqlab KAPA Library Quantification Kit and the Applied Biosystems 7900HT Sequence Detection System. The pool was sequenced on an Illumina NovaSeq6000 sequencing instrument with a PE100 protocol.

### Analysis of RNAseq

RNA-seq was performed with a directional protocol. Quality control, trimming, and alignment were performed using the nf-core (Ewels et al., 2020). RNA-seq pipeline (v3.0) (doi.org/10.5281/ZENODO.1400710). Details of the software and dependencies for this pipeline can be found at the referenced DOI for the pipeline and: https://github.com/nf-core/rnaseq/blob/master/CITATIONS.md. The reference genome sequence and transcript annotation used were *Homo sapiens* genome GRCh38 from Ensembl version 103.

Differential expression analysis was performed in R version 4.1.2 (2021-11-01) (R Core Team 2021) with DESeq2 v1.34.0 (Love et al., 2014) to make pairwise comparisons between groups. Log Fold Change shrinkage estimation was performed with ashr (Stephens, 2016). Only genes with a minimum coverage of 10 reads in all samples from each pairwise comparison were considered as candidates to be differentially expressed. Genes were considered deferentially expressed if they showed a *|log2(Fold Change)| > 1* and were below a FDR of 0.05. Genes with a minimum coverage of 10 reads in all samples from each pairwise comparison were included in functional enrichment analyses, and considered as the ‘gene universe’ for over-representation based analyses. Functional enrichment analysis was performed with clusterProfiler v4.2.0 (Yu et al., 2012).

### Immunofluorescence staining

Cells were washed once with PBS 1X (Thermo Fisher, #18912014) and fixed at RT for 15 min in 4 % PFA pH 7.4 (Sigma-Aldrich, #P6148-500G). After washing with PBS, cells were incubated for 10 min at RT in PBS containing 0.2 % Triton X-100 (Fluka, #93418) for permeabilization. Subsequently, cells were washed again and treated with 10 % goat serum in PBS for 10 min at RT. Depending on the primary antibody, cells were incubated for 2 hours with the antibody diluted in 1% goat serum in PBS. After three washing steps with PBS for 5 min, cells were incubated for 1 hour in a secondary antibody diluted in 1 % goat serum in PBS. Cells were washed again three times for 5 min with PBS, while DAPI (Sigma-Aldrich, #D9542-1MG) was added during the second wash. Samples were mounted with FluorSave reagent (Millipore, #345789). The antibodies used in this study are the following: anti-ERLIN2 (1:1000, Cell Signaling, 2959), anti-TMUB1 (1:1000, Abcam, ab180586), anti-GM130 (1:1000, BD Biosciences, 610822), Alexa Fluor 488 anti-rabbit IgG (1:1000, ThermoFisher, # A-11008) and Alexa Fluor 594 anti-mouse IgG (1:1000, ThermoFisher, # A-11012).

For LDs staining no permeabilization was performed and Bodipy 493/503 (ThermoFisher, D3922) was applied just after fixation with PFA 4% for 30 min at final concentration of 5 µM and washed five times for 10 min in PBS.

### Transfection and Live imaging

Transfection of cells was performed using lipofectamine 2000 (Invitrogen; #11668019), 24h after seeding. Before the transfection, medium of the cells was changed to OptiMEM. For the transfection, 2 µl of lipofectamine were diluted in 250 µl of OptiMEM (Gibco, #11058021) and kept for 5 min at room temperature. Afterwards, the Lipofectamine:OptiMEM mix was added to 250 µl of OptiMEM containing 1ug of DNA. After 15 min the DNA:Lipofectamine complex was added dropwise to the cells and after 6h incubation, the medium was exchanged by DMEM. Live imaging was performed 24h after the transfection and at 37°C with 5% CO_2_ to resemble the incubator conditions. The RUSH system was used to analyse ER to Golgi trafficking similarly as described before (PubMed 22406856). Cells were transfected 24h before the imaging with Str-Ii_VSVG-SBP-EGFP (Addgene #65300). Live imaging was performed with a spinning disk microscope for 45 min and one frame was acquired every 30 sec. Acquisition of the video was initiated directly after the biotin was added to the medium.

### Image analysis

Image analysis was performed using Fiji/ImageJ 2.9.0 (Schindelin et al., 2012).

ER morphology was quantified similarly to Hu et al (Hu et al., 2009), i.e., by visual inspection cells were classified into three categories (normal, intermediate or collapsed) according to ER shape using RTN4b immunostaining. The RTN4b area was calculated using the same staining after a Gaussian blur (Sigma = 20) filter was applied. The area was later obtained by thresholding with fixed values for each biological replicate.

For the quantification of the Golgi apparatus and LDs, after manual segmentation of the cells, the Golgi apparatus and LDs were automatically segmented using GM130 immunostaining and Bodipy staining, respectively. Segmentation was performed using a fixed threshold for all genotypes in each biological replicate. The objects derived by thresholding were counted per cell using the “Analyze particles” function available in Fiji.

For VSVG-GFP/GM130 colocalization, after subtraction of the background calculated from an area outside the cells, the cells were manually segmented and for each cell both the total VSVG-GFP fluorescence intensity and the fluorescence intensity within a mask obtained from GM130 signal thresholding were calculated. The value used is the ratio between the fluorescence intensity inside the Golgi and the total fluorescence intensity for each cell. Scalebar is equal to 10 µM.

### Protein extraction and Immunoblotting

Cells were harvested by scarping and the pellet was lysed in an equal volume of RIPA buffer (50 mM TRIS-HCl pH 7.4, 150 mM NaCl, 1 mM EDTA, 1 % Triton X-100, 0.25 % sodium deoxycholate, 1mM sodium orthovanadate, Protease Inhibitor Cocktail powder (Sigma, # P2714-1BTL)) for 30 min on ice. Samples were centrifuged for 30 min at 20,000 x *g* at 4 °C and protein concentrations of the supernatants were measured with standard Bradford assay (Bio-Rad Laboratories). 25 or 50 µg of proteins were diluted in Laemmli buffer 3× (188 mM Tris-HCl, 6 %SDS, 12 % Glycerol, 0.05 % Bromophenolblue, 10% β-Mercaptoethanol) and then analysed by SDS-PAGE followed by immunoblot analysis. Stacking gels (1% w/v Ammonium persulfate, 0.1 % SDS, 0.1 % N,N,N’,N’-Tetramethylethylendiamin and 0.125 M Tris-HCl pH 6.8) were casted with a final concentration of 6% of acrylamide. Resolving gels (1% w/v Ammonium persulfate, 0.1 % SDS, 0.1 % N,N,N’,N’-Tetramethylethylendiamin and 0.375 M Tris-HCl pH 8.8) were casted at percentages of 15%, 12% or 10% of acrylamide depending on the target protein. Protein separation was performed in running buffer (25 mM Tris, 192 mM glycine, 0.1 % SDS) for up to 2 h and at 50-100 V depending on the gel concentration and the molecular weight of the protein of interest. Proteins were blotted on polyvinylidene difluoride membrane (PVDF) (Amersham, #10600023) of pore size 0.45 µm using wet chambers and blotting buffer (25 mM Tris, 192 mM glycine, 20 % methanol) at 4°C for either 1.5h at 300 mA or for 16 h at 30 V. After blotting, proteins were stained for 10 min in PonceauS and a picture of the membrane was acquired. Then membranes were blocked for 30 min in skim milk (Carl Roth, #P2714-1BTL) 5% in TBST (20 mM Tris-HCl pH 7.4, 150 mM NaCl, 0.1 % Tween-20) at room temperature. Primary antibodies were diluted in the same blocking solution and incubated over night at 4°C or at room temperature for 2h. Respective HRP-linked secondary antibodies were used at 1:10,000 dilution for both anti-mouse and anti-rabbit primary antibodies and incubated for 1h at room temperature. After incubation with both primary and secondary antibody membranes were washed three times in TBST. HRP was detected using ECL western blotting detection reagents (VWR, # RPN2106). Chemiluminescence was detected using and X-ray films (Bema, # 4005194) or the Vilber Fusion Solo S. Intensities of bands of interest were quantified using ImageJ.

The antibody used in this study are the following: anti-ERLIN1 (1:1000, Sigma, (HPA011252)), anti-ERLIN2 (1:1000, Cell Signaling, (2959)), anti-TMUB1 (1:1000, Abcam, (ab180586)), anti-TMUB2 (1:1000, Proteintech, (28044-1-AP)), anti-GAPDH (1:2000, Millipore, #MAB374), anti-Flotillin1 (1:1000, BD Transduction Laboratories, #610821), anti-KIDINS220 (1:1000, Thermo Fisher, PA5-22116), anti-RTN4b (1:1000, Novus Biologicals, NB100-56681), anti-VDAC2 (1:1000, CellSignaling, 9412S), anti-TMEM259 (1:1000, AtlasAntibodies, HPA042669), anti-PMP70 (1:1000, Abcam, ab85550), anti-Flag (1:5000, Sigma-Aldrich, # F7425), HRP-linked anti-mouse IgG (1:10000, Sigma-Aldrich, #A9044), HRP-linked anti-rabbit IgG (1:10000, Sigma-Aldrich, #A0545).

### Immunoprecipitation

For IPs, 1.5×10^7^ cells seeded on a 15 cm dish one day before the lysis. The next day, cells were washed once with PBS at room temperature, harvested with a cell scraper, pelleted at 60 x *g* for 3 min and resuspended in 400 µl IP buffer (Tris-HCl pH 7,4 50 mM, KCl 50 mM, Triton X-100 0,5%, Protease Inhibitor Cocktail powder (Sigma, # P2714-1BTL)). Then, cells were passed six times through a 29 Gauge needle, incubated for 30 min on ice and centrifuged at 20,000 g for 30 min at 4°C. Protein concentration was determined via a Bradford assay, 500 µg of proteins were transferred in a new microcentrifuge tube and the volume adjusted to 250 µl using IP buffer. Next, 0.5 µg of the respective antibody was added to the proteins and incubated overnight at 4 °C on a rotor. An anti-RFP (Rockland, 600-401-379S) was used a control antibody for the TMUB1 IP. Afterwards, 20 µl of Dynabeads™ Protein G (Thermofisher, # 10003D) were equilibrated three times with 1 ml of IP buffer using a magnetic rack, added to the protein:antibody lysate and incubated for 3 h at 4 °C. After incubation, the magnetic rack was used to separate the beads from the flow-through and beads were washed three times with 1 ml of IP buffer. Depending on the followed application, elution was performed in either 30 µl of Laemmli buffer 3X, for western blot analysis, or in SP3 buffer (5% SDS in 1x PBS) for proteomics analysis. After addition of the respective elution buffer, beads were vortexed for 1 min and incubated at 95 °C for five min. The eluate was separated from the beads using the magnetic rack.

### Detergent resistant membrane isolation

DRMs were isolated according to the protocol from George and colleagues (George et al., 2010) with following changes. Lipid-protein crosslinking step was not performed and the Triton X-100 concentration of the isolation buffer was increased to 0.5 %. Per sample, 1.5X10^7^ cells were seeded on 15 cm plates and after two days washed twice with cold PBS + 5 mM EDTA, harvested using a cell scraper and centrifuged for 10 min at 11.200 x *g* and at 4 °C. Cell pellets were kept overnight at –80 °C. Samples were washed once with TBS and lysed in raft isolation buffer (0.5% Triton X-100, 150 mM NaCl, 5 mM EDTA, 50 mM TRIS-HCl pH 7.4). Following lysis, cell lysates were centrifuged for 15 min at 112 x *g*, 4 °C and protein concentrations were determined via Bradford assay. 1 mg of proteins were diluted in 500 μl raft isolation buffer and subsequently mixed with 1 ml 60 % Opti-prep (Axis-shield, # 1114542) solution to obtain a final concentration of 40 % Iodixanol. Homogenates were transferred into ultra-clear tubes (Beckman and Coulter, # 344062) and overlaid with a step gradient of 30 % iodixanol solution and 5 % iodixanol solution to a final volume of 4 ml. The step gradients were centrifuged for 5 h at 132.000 x *g* at 4 °C (SW60 Beckman and Coulter, # 335649). Subsequently, five fractions were obtained by taking 800 μl from the top to bottom.

### Proteomic analysis of co-IP’s, post nuclear lysate and SILAC-labelled DRMs

#### Sample preparation

After IP, the proteins eluted in SP3 buffer were reduced, alkylated and digested with trypsin using the SP3 protocol originally described by Hughes et. al (Hughes et al., 2019). For the analysis of DRM fractions generated from SILAC labelled cells and label-free profiling of post-nuclear lysate, the proteins were precipitated with four times the volume of ice-cold acetone by incubation at –80°C for 15 minutes followed by 2h at –20°C. After centrifugation at 16,000 g for 15 minutes at 4°C, the supernatant was discarded and the pellet was washed with 500 μl of ice-cold acetone and centrifuged twice for 5 min at 16,000 g. The air-dried pellet was resuspended in 8 M urea in 50 mM triethylammonium bicarbonate (TEAB) pH 8.5 and digested with LysC and Trypsin as described previously (Schatton et al., 2022). Digested samples were loaded on Stage Tips and submitted to the CECAD proteomics facility.

#### Data acquisition

Immunoprecipitated as well as SILAC-labelled samples were analysed in data dependent acquisition mode (DDA) on a Q-Exactive Plus (Thermo Scientific) mass spectrometer) as described previously (Schatton et al., 2022). For whole proteome profiling of post-nuclear lysate, samples were analysed in data independent acquisition mode (DIA) on an Orbitrap Q-Exactive plus (Thermo Scientific) mass spectrometer that was coupled to an easy 1000 nano lc system (Thermo Scientific). Samples were loaded onto an in-house packed analytical column (50 cm — 75 µm I.D., filled with 2.7 µm Poroshell EC120 C18, Agilent) equilibrated in buffer A (0.1 % formic acid in water). Peptides were chromatographically separated with a flow rate of 250 nL/min and a 110 min gradient, followed by a 10 min column wash with 95% solvent B (0.1 % formic acid in 80% acetonitrile. To generate an experiment-specific gas phase fractionated library, a pool of all samples was used for narrow window DIA measurements covering the range from 400 m/z to 1000 m/z (Searle et al., 2020) with six consecutive 100 m/z acquisitions. MS1 scans of the 100 m/z gas phase fraction were acquired at 30k resolution. Maximum injection time was set to 50 msec and the AGC target to 3E6. MS2 scans were acquired in 25 x 4 m/z staggered windows resulting in 50 nominal 2 m/z windows after demultiplexing. MS2 settings were 17,500 resolution, 60 msec maximum injection time and an AGC target of 1E6. For the acquisition of the sample data, an identical chromatography method was used. MS1 scans were acquired at 35,000 resolution, maximum injection time was set to 60 msec and the AGC target to 1E6. DIA scans covering the precursor range from 400 – 1000 m/z and were acquired in 25 x 25 m/z staggered windows. MS2 spectra starting at 200 m/z were acquired at 17,500 resolution with a maximum injection time of 60 msec and an AGC target of 1E6.

#### Data processing

DDA raw data were processed with MaxQuant version 1.5.3.8 or 2.0.1.0 (Cox et al., 2008, Cox et al., 2011) as described previously (Schatton et al., 2022). LFQ quantification was enabled with default settings and the match-between runs option was enabled within replicate groups. For SILAC ratio quantification the label multiplicity was set to 3 with Arg-6 and Lys-4 selected as medium and Arg-10 and Lys-8 as heavy labels. DIA data were analysed with DIA-NN 1.8.1 (Demichev 2020). First the narrow window DIA runs were demultiplexed and transformed to mzML files using the msconvert module in ProteoWizard. These files were used to generate an experiment specific spectral library from a SwissProt human canonical fasta file using the library generation option in DIA-NN. The settings differing from default were: Min precursor m/z set to 400, Max precursor m/z set to 1000, Heuristic protein inference and no shared spectra. The experiment specific spectral library (10471 protein isoforms, 11109 protein groups and 104704 precursors) was used for subsequent sample analysis. For statistical analysis the MaxQuant or DIA-NN output tables were imported to Perseus version 1.6.15.0 (Tyanova et al., 2016). Potential contaminants were removed, intensities were log2 transformed and the data sets were filtered for at least 100 % data completeness in at least one condition. Remaining missing values were imputed with random values from the lower end of the intensity distribution using Perseus defaults. Student’s t-tests and ANOVAs were calculated with permutation-based FDR control (q < 0.05). For the five DRM fractions, no imputation was use and the following steps were performed separately. Significant candidates with changing SILAC ratios between any conditions were identified by ANOVA multiple sample testing (S0 = 0.1, permutation-based FDR < 0.05 with 250 randomizations). Next, the dataset was filtered for significantly regulated candidates, SILAC ratios were Z-score normalized protein-wise, and Euclidean hierarchical clustering was performed to identify protein clusters with similar changing protein SILAC ratios over the three conditions. Finally, all fractions were merged, protein groups were annotated with Gene ontology (GO) terms and compartments (Itzhak et al., 2016). and Fisher’s exact tests were performed for each fraction separately (Benjamini-Hochberg FDR = 0.02) to identify regulated processes systematically.

### Lipidomics

#### Triacylglycerols and cholesteryl esters

Cells were homogenized in Milli-Q water (approx. 10^6^ cells/100 µl) using the Precellys 24 Homogenizer (Peqlab) at 6,500 rpm for 30 sec. The protein content of the homogenate was routinely determined using bicinchoninic acid.

To 50 µl of cell homogenate 450 µl of Milli-Q water, 1.875 ml of chloroform/methanol/37 % hydrochloric acid 5:10:0.15 (v/v) and internal standards (30 µl of d5-TG Internal Standard Mixture I and 256 pmol cholesteryl ester 19:0; Avanti Polar Lipids) were added. Lipid extraction was performed as previously described (Kumar et al., 2015).

Triacylglycerols and cholesteryl esters were analysed by Nano-Electrospray Ionization Tandem Mass Spectrometry (Nano-ESI-MS/MS) with direct infusion of the lipid extract (Shotgun Lipidomics) as previously described (Hammerschmidt et al., 2019; Ozbalci et al., 2013). Endogenous lipid species were quantified by referring their peak areas to those of the respective internal standard. The calculated lipid amounts were normalized to the protein content of the cell homogenate.

#### Cholesterol

Cholesterol levels in cells were determined by Liquid Chromatography coupled to Electrospray Ionization Tandem Mass Spectrometry (LC-ESI-MS/MS): To 30 µl of the cell homogenate mentioned above, 70 µl of Milli-Q water and 1.26 nmol d7-cholesterol as internal standard (Avanti Polar Lipids) were added. Lipid extraction and LC-MS/MS analysis of cholesterol were performed as previously described (Mourier et al., 2015).

#### Ceramides and sphingomyelins

Levels of ceramides and sphingomyelins in isolated DRMs were determined by LC-M/MS by treating the samples using a procedure previously described (Ejsing et al., 2009) with some modifications:

To 70 µl of DRM preparation 130 µl of 155 mM ammonium carbonate buffer were added. Lipids were extracted by adding 990 µl of chloroform/methanol 17:1 (v/v) and internal standards (127 pmol ceramide 12:0 and 124 pmol sphingomyelin 12:0; Avanti Polar Lipids), followed by shaking at 900 rpm/min in a ThermoMixer (Eppendorf) at 20 °C for 30 min. After centrifugation (12,000×g, 5 min, 4 °C), the lower (organic) phase was transferred to a new tube, and the upper phase was extracted again with 990 mL chloroform/methanol 2:1 (v/v). The combined organic phases were dried under a stream of nitrogen. The residues were resolved in 100 µL of Milli-Q water and 750 µL of chloroform/methanol 1:2 (v/v). Alkaline hydrolysis of glycerolipids and LC-ESI-MS/MS analysis of ceramides and sphingomyelins were done as previously published (Oteng et al., 2017).

### AlphaFold Multimer prediction of ERLIN1, ERLIN2, TMUB1 and RNF170 complex

Multimer predictions of the ERLIN1-ERLIN2-TMUB1-RNF170 complex were performed using the Cologne High Efficient Operating Platform for Science (CHEOPS) provided by the Regional Computing Center of Cologne (RRZK) utilizing 4 CPU cores of a node with 2x Intel(R) Xeon(R) Gold 6248 CPU @ 2.50GHz, 60 GB RAM and a Nvidia Tesla V100-SXM2 GPU with 32 GB VRAM running the AlphaFold module 2.2.0 according to (Jumper et al., 2021), with 5 seeds per model generating in total 25 models. Peptide sequences of ERLIN1 (sp|O75477|ERLN1_HUMAN Erlin-1), ERLIN2 (sp|O94905|ERLN2_HUMAN Erlin-2), TMUB1 (sp|Q9BVT8|TMUB1_HUMAN Transmembrane and ubiquitin-like domain-containing protein 1),RNF170 (sp|Q96K19|RN170_HUMAN E3 ubiquitin-protein ligase RNF170), VCP (sp| P55072|TERA_HUMAN Transitional endoplasmic reticulum ATPase) and FAF2 (sp|Q96CS3|FAF2_HUMAN FAS-associated factor 2) were extracted from Uniprot.org. The top-ranked model, with the highest confidence, of the Multimer prediction was used for further analysis. AlphaFold models were visualized using the latest version of ChimeraX-1.5 and hydrogen bonds were predicted using the ChimeraX function “H-Bonds”.

### Plasmids

Str-Ii_VSVG-SBP-EGFP was a gift from Franck Perez (Addgene plasmid # 65300; http://n2t.net/addgene:65300; RRID:Addgene_65300).

pBABE-puro was a gift from Hartmut Land & Jay Morgenstern & Bob Weinberg (Addgene plasmid # 1764; http://n2t.net/addgene:1764; RRID:Addgene_1764)

pERLIN2-3xFlag, pERLIN2-3xFlag (R36K), pERLIN2-3xFlag (G48V), pERLIN2-3xFlag (H50Y), pERLIN2-3xFlag (R36K-H50Y) and pERLIN2-3xFlag (R36K-G48V-H50Y) were generated from the p3XFLAG-CMV-14. All the mutations were tested via Sanger sequencing.

For the Cloning, ERLIN2 was amplified with 5’-TTAGAATTCATGGCTCAGTTGGGAGCAGTT-3’ and 5’-AATGGATCCATTCTCCTTAGTGGCCGTCTC-3’ from cDNA and the product was cloned into into p3XFLAG-CMV-14 via restriction enzyme cloning using EcoRI and BamHI. The R36K variant was generated with the Q5® Site-Directed Mutagenesis Kit (NEB, # E0554S) from the pERLIN2-3xFlag using the manufacturer’s protocol (primers sequence: Fwd 5’-GTATATTACAaAGGCGGTGCC-3’ and Rev 5’-CCCAATATGTCCCTCTTCTATC-3’). All the other variants were generated using the Gibson assembly (NEBuilder® HiFi DNA Assembly Master Mix, # E2621S) from the pERLIN2-3xFlag using the manufacturer’s protocol. The pERLIN2-3xFlag was amplified using 4 primers, 2 binding the backbone, common for all the mutations (Fwd 5’-TAATGAGTGAGCTAACTCACATTAATTGCGTTGCG-3’ and Rev 5’-GTGAGTTAGCTCACTCATTAGGCACCCC-3’), and other 2 primers specific for the desired mutation (G48V: 5’-CAGCGGCCCTGtTTTCCATCTCATGCTCC-3’ and 5’-GATGGAAAaCAGGGCCGCTGGTCGAAGTC-3’; H50Y: 5’-CCCTGGTTTCtATCTCATGCTCCCTTTCATC-3’ and 5’-GCATGAGATaGAAACCAGGGCCGCTGGTC-3’; G48V-H50Y: 5’-CAGCGGCCCTGtTTTCtATCTCATGCTCCCTTTC-3’ and 5’-GATaGAAAaCAGGGCCGCTGGTCGAAGTC-3’).

### siRNA

For TMUB1 downregulation stealth siRNA from Thermo Fisher were used at 100 nM final concentration. After two days from the first transfection, performed using lipofectamine 2000 like previously described, a second transfection was performed and the cells were analysed after two days. Shorter protocols failed to downregulate TMUB1 in our system. Three siRNA were used: 5’-ACCUCCCUCCCAACUGCGUUCUCCA-3’, 5’-GGGAACAGCAGGUGCGACUCAUCUA-3’ and 5’-GCCAUGGCAGCUACCGACAGCAUGA-3’.

### Statistical analysis

A one-way analysis of variance followed by a post-hoc Tukey’s test was used to compare multiple groups. The statistical analysis of the mass spectrometry analysis is described in detail in the “Data processing” section of the proteomics section.

## Supplemental material

Figure S1 shows the distribution of selected ERLIN2 interactors. Figure S2 shows the TMUB1-L KO cell line and the specificity of the TMUB1 antibody. Figure S3 shows ERLINs depletion affects adhesion pathway and cell migration. Figure S4 displays the intensity of selected proteins in different fractions measured by SILAC. Figure S5 shows trafficking experiments using the RUSH system. Table S1 contains results of proteomics experiments. Table S2 shows results of transcriptomics. Table S3 shows SILAC experiments. Table S4 contains lipidomics data. Videos 1-3 show live imaging RUSH assays in WT, DKO and DKO^+E1/2^.

## Data availability statement

All data supporting the findings of this study are available within the paper and its Supplementary Information. Proteomics raw data are available via ProteomeXchange with identifier PXD048503.

## Acknowledgments

The authors thank Esther Barth, the CECAD proteomics, lipidomics, imaging and FACS facility for support and members of the Rugarli laboratory for constructive discussions. This work was funded by the Deutsche Forschungsgemeinschaft (RU 1653/4-1) and by the Walter and Monika Neupert foundation to M.V. E.I.R. and M.V. are members of the RTG-Reloc (411422114-GRK 2550) funded by the DFG. The authors declare no competing financial interests.

## Author Contribution

M. Veronese: conceptualization, formal analysis, investigation, visualization, writing, review and editing. S. Kallabis: SILAC proteomics analysis, visualization, writing, review and editing. A. Kaczmarek: AlphaFold analysis, visualization, writing, review and editing. L. Roberts: investigation, visualization, and review. A. Das: investigation, visualization, review and editing. S. Schumacher: investigation. A. Lofrano: investigation. S. Brodesser: formal analysis, investigation, writing, review and editing. S. Müller: formal analysis, investigation, writing, review and editing. K. Hofmann: formal analysis. M. Krüger: conceptualization, supervision, review and editing. E.I. Rugarli: conceptualization, supervision, funding acquisition, writing original draft, review, and editing.

## Supplementary Figures

**Figure S1.**
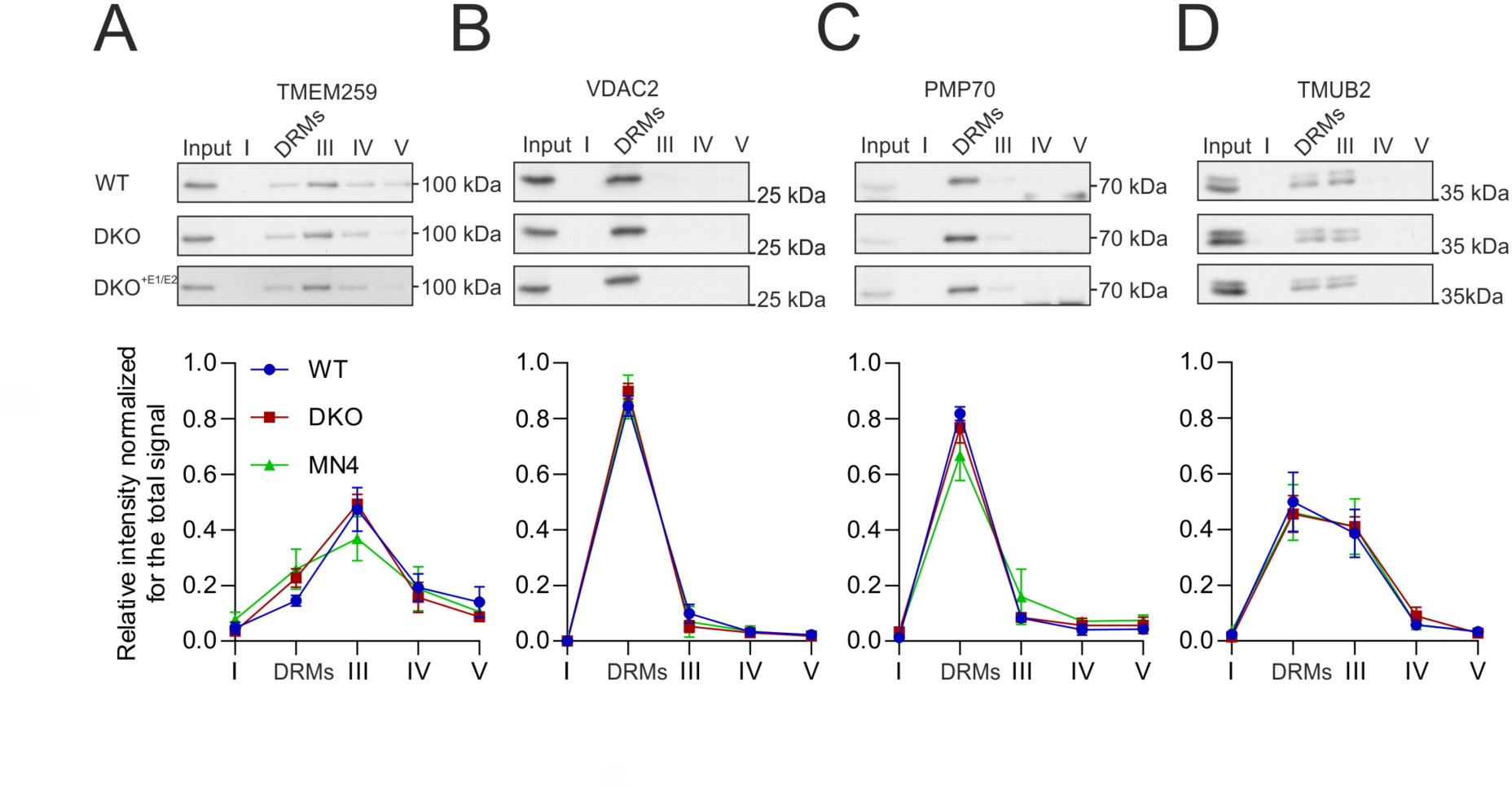
Selected ERLIN2 interactors do not change distribution on a DRM-isolating gradient in DKO cells. Western blot of TMEM259 **(A)**, VDAC2 **(B)**, PMP70 **(C)** and TMUB2 **(D)** with respective quantification of the distribution in different fractions. None of the proteins show significant difference using one-way analysis of variance with post Tukey’s multiple comparison test.

**Figure S2.**
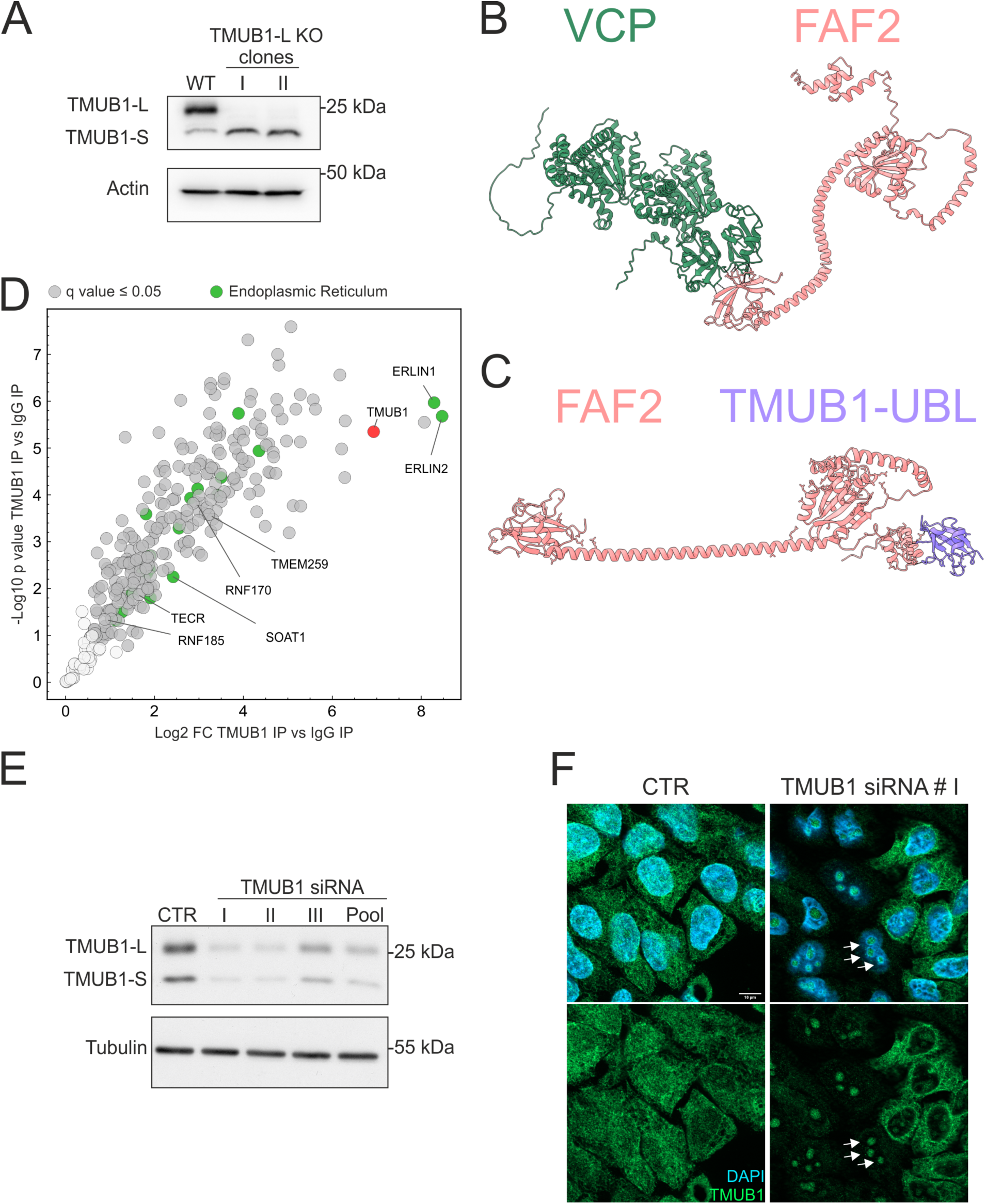
TMUB1-L KO cell line and specificity of the TMUB1 antibody. (**A**) Western blot of two knock-out clones for TMUB1-L. Actin is used as loading control. **(B)** AlphaFold Multimer model of VCP-FAF2. **(C)** AlphaFold Multimer model of FAF2-TMUB1-L-UBL. **(D)** Volcano plot showing enriched proteins by a TMUB1 antibody versus control IgG. Proteins significantly enriched by TMUB1 (q value ≤0.05) are labelled in dark grey and proteins belonging to the GOCC term endoplasmic reticulum are labelled in green. N= 4 biological replicates. **(E)** Western blot of HeLa cells treated with different siRNA against TMUB1. **(F)** TMUB1 immunofluorescence in HeLa cells after downregulation using the siRNA number I. Scalebar = 10 µm.

**Figure S3.**
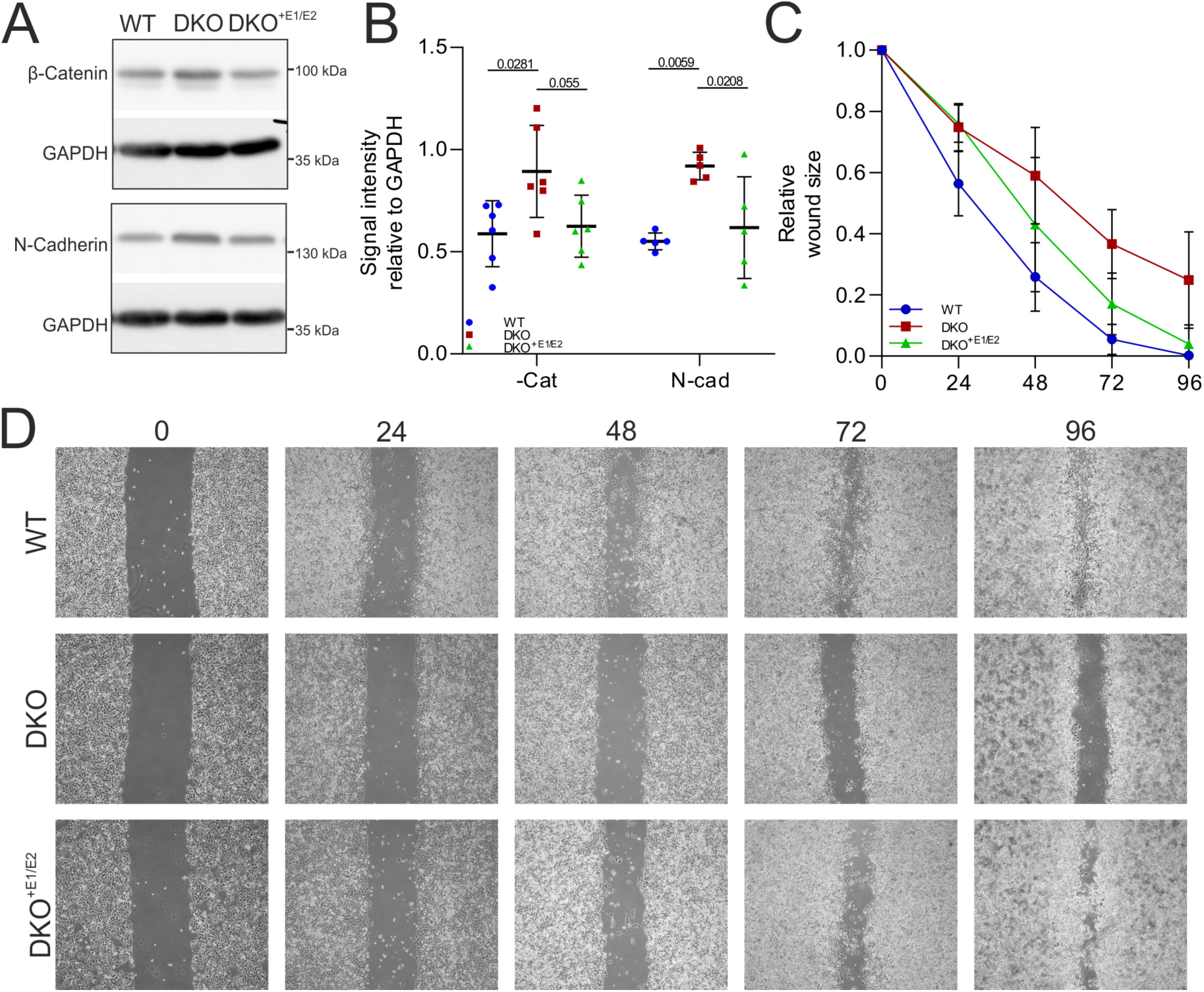
ERLINs depletion affects adhesion proteins and cell migration. (**A, B**) Western blot in WT, DKO and DKO^+E1/E2^ for β-Catenin and N-Cadherin with quantification. Dots are biological replicates. **(C, D)** Quantification (C) and representative images (D) of a scratch assay in WT, DKO and DKO^+E1/E2^.

**Figure S4.**
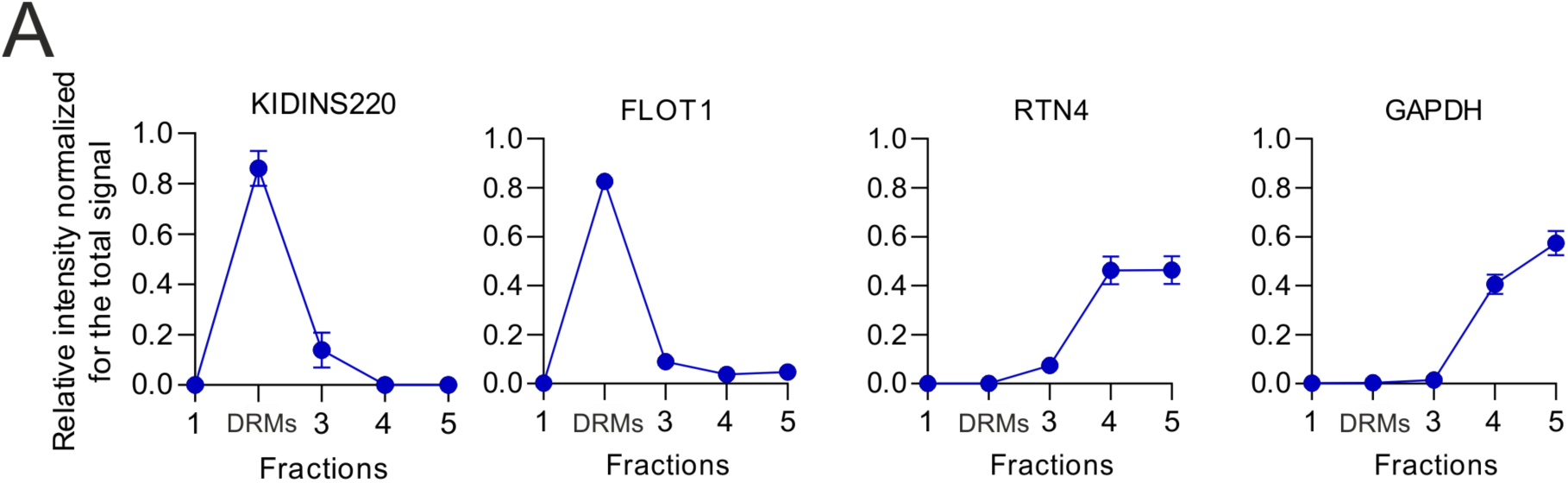
Intensity of selected proteins in different fractions measured by SILAC (A) Purity of the SILAC DRMs isolation was tested using the localization of FLOT1 and KIDINS220 as DRM markers, RTN4 as a non-DRM membrane protein and GAPDH as soluble protein.

**Figure S5.**
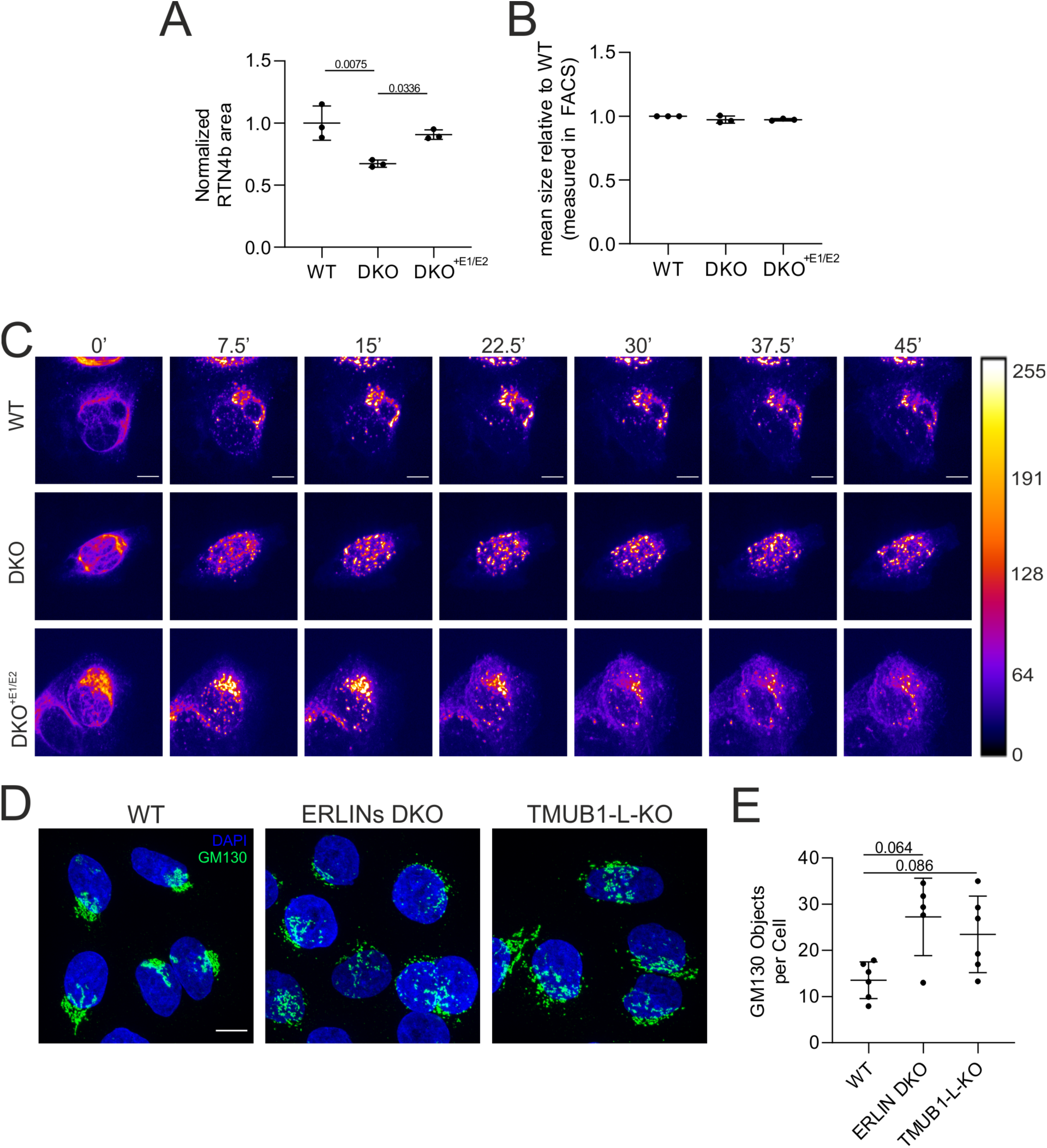
DKO cells display defects in post-Golgi trafficking of VSVG-GFP. (**A**) Quantification of the area occupied by the RTN4 staining in cells of different genotypes. **(B)** Cell size measured in cytofluorimetry. **(C)** Live imaging RUSH assay using the VSVG-GFP construct, pictures show different timepoints of the video. A Fire LUT was applied using ImageJ in order to show the levels of fluorophore accumulating in the Golgi apparatus. The bar represents the value of the pixel. **(D)** GM130 immunofluorescence of WT, ERLIN DKO and TMUB1-L KO cells. Scalebar = 10 µm. **(E)** Quantification of Golgi fragmentation. N= 5 biological replicates (at least 200 cells were analysed per genotype). Statistical test shown is the paired one-way analysis of variance with post Tukey’s multiple comparison test.

